# A bibliometric analysis of research on the mitochondrial roles in prostate cancer and the virtual design of LONP1 - specific antibodies using the GeoBiologics platform

**DOI:** 10.1101/2025.03.14.643215

**Authors:** Jiawei Pan, Yuan Zhang, Linglong Jiang, Yuwei Shen, Yangyang Sun, Jundong Zhu, Zhen Chen, Min Fan, Jian Shi

## Abstract

**Background:** Prostate cancer remains one of the most prevalent malignancies among men globally, with its incidence showing an upward trend worldwide. Mitochondria, as central regulators of cellular energy metabolism, play crucial roles in prostate cancer initiation, progression, and drug resistance mechanisms. While mitochondria-targeted therapeutic strategies have emerged as a significant focus in cancer research in recent years, comprehensive bibliometric analyses mapping the evolving landscape of this field remain scarce. This study systematically investigates research trends in mitochondrial-prostate cancer interactions through bibliometric methods, identifying LONP1 as an emerging research focus in mitochondria-related prostate cancer therapy. Building on these findings, we employed artificial intelligence to virtually design a LONP1-specific antibody, proposing novel therapeutic targeting strategies for this field.

**Methods:** Utilizing the Web of Science Core Collection database (2015-2023), we conducted visualization analyses through CiteSpace and VOSviewer to map network relationships among countries, institutions, journals, authors, and keywords. Building on this foundation, a humanized antibody targeting LONP1 was computationally designed and screened through the GeoBiologics platform.

**Results:** Analysis of 452 included publications revealed the United States and China as leading contributors in this research domain. The field has progressively transitioned from fundamental mechanistic investigations to clinical applications, particularly focusing on drug resistance mechanisms, and combination therapy. LONP1 was identified as a critical mitochondrial regulator strongly associated with prostate cancer progression. Our AI-designed antibody (Antibody_82) demonstrated superior binding affinity and stability through effective targeting of LONP1’s ATP-binding site.

**Conclusion:** This bibliometric study delineates evolving research trends in mitochondrial involvement in prostate cancer. The developed LONP1-targeting antibody shows promising therapeutic potential for castration-resistant prostate cancer (CRPC) patients, potentially offering more effective treatment alternatives.

## 1. Introduction

Prostate cancer is one of the most prevalent malignant tumors in men worldwide, with significant variations in its incidence and mortality rates across different regions and ethnic groups[1]. As the global population ages and lifestyle patterns evolve, the incidence of prostate cancer continues to increase [2].

Mitochondria serve as the primary sites of cellular energy production, playing a crucial role in energy metabolism and being involved in essential biological processes, such as cell signaling, cell cycle regulation, and apoptosis[3]. Accumulating evidence has underscored the strong association between mitochondrial dysfunction and the onset and progression of various diseases. This connection is particularly significant in cancer biology, where the role of mitochondria has attracted considerable attention in prostate cancer research, studies have demonstrated that mitochondria are involved in multiple facets of tumor initiation, progression, and drug resistance. Specifically, mutations in mitochondrial DNA, alterations in mitochondrial metabolic pathways, and disruptions in mitochondrial dynamics are closely linked to the progression of prostate cancer[4,5]. Moreover, mitochondrial-targeted therapeutic strategies, such as promoting mitochondria-mediated apoptosis, have emerged as promising approaches for the treatment of prostate cancer[6].

Bibliometric analysis is a research method that quantifies and analyzes literature data to reveal the current state and trends of a specific field of study[7]. This study aims to apply bibliometric methods to systematically analyze research literature related to mitochondria and prostate cancer, exploring the research hotspots, development trends, and potential academic frontiers in this field. By retrieving relevant literature on mitochondria and prostate cancer from the Web of Science database, we collected studies published between 2015 and 2023. Using visualization analysis tools such as Citespace and VOSviewer, we identified the evolution and frontier dynamics of research in this area. Additionally, we summarized relevant experimental information, including experimental methods and treatment protocols. By analyzing genes associated with mitochondrial function that are closely linked to the progression of prostate cancer, it has been identified that LONP1 is an emerging research hotspot. Therefore, we virtually designed specific antibodies against LONP1 using the GeoBiologics platform, an AI-powered one-stop platform for antibody design and optimization.

## 2. Materials and methods

### 2.1. Data collection and pre-processing

This study utilized data from the Science Citation Index Expanded (SCIE) within the Web of Science Core Collection (WoSCC). To ensure search accuracy, the PRISMA guidelines for literature collection were followed. The search formula was constructed as follows:TS = (“mitochondria” OR “mitophagy”) AND (“prostatic cancer” OR “prostatic carcinoma” OR “prostatic adenocarcinoma” OR “prostate tumor” OR “prostatic malignancy” OR “prostatic neoplasm” OR “radical prostatectomy”), with language restricted to “English,” document types including “dissertation,” “online publication,” and “review paper,” and a time frame spanning from 2015 to 2023. Ultimately, 452 relevant articles were retrieved.

During the data preprocessing phase, the following steps were implemented in this study: First, the researchers exported the complete records of all retrieved articles and their citations in plain text format. Second, key elements of the articles (such as year, citation frequency, title, author keywords, additional keywords, journal, authors, institutions, and countries) were extracted into an Excel file. The third step involved removing all duplicate article records. Fourth, missing elements in the articles were supplemented. Fifth, irrelevant terms were filtered out, and synonyms were merged.

Lastly, the standardization of element formatting was achieved by eliminating unnecessary spaces and punctuation marks.

### 2.2. Analysis and visualization

In this study, we primarily used VOSviewer (version 1.6.16), Citespace (version 5.8.R2), and the Bibliometric Online Analysis Platform (http://bibliometric.com/) to identify network features such as co-cited articles, keywords, countries, institutions, journals, authors, and keywords, and to visually present the research findings. Co-authorship analysis, co-citation analysis, and co-occurrence analysis are common methods in bibliometric analysis. Co-authorship analysis explores the collaborative relationships between authors, countries, or institutions by analyzing the number of jointly authored papers. Co-occurrence analysis quantitatively assesses the relationships between different items by measuring whether they appear together. Co-citation analysis shows the strength of relationships between cited items based on the number of citations[8].

VOSviewer was mainly used for citation/co-citation analysis of countries and institutions, as well as keyword co-occurrence analysis. Additionally, we employed Citespace for co-authorship analysis of literature and keywords. Citespace also generated dual-map overlays to visually present the evolutionary trends within the field. Furthermore, we utilized the online bibliometric analysis platform for co-authorship analysis by country and publication analysis.

## 3. Results

### 3.1. The annual trend of paper publication quantity and citations

Through meticulous searching and screening, we collected a total of 452 research papers related to mitochondria and prostate cancer. These articles have made a significant impact in the academic community, with a cumulative total of 17,774 citations, highlighting the importance and attention of this research field. In Figure 1, over the nine-year period from 2015 to 2023, the number of publications significantly increased between 2013 and 2017, followed by a decline after 2018. Specifically, from 2018 to 2020, the number of publications stabilized; there was an increase in 2021, but a decline was observed again in 2023. The total number of citations for all articles was 19,406, with 19,132 citations excluding self-citations. The average citation per article was 42.32, and the H-index was 59. The H-index, which takes both the number of publications and citations into account, is a key indicator of research impact. A higher H-index reflects the quality and influence of research in the field.

**Figure 1.**
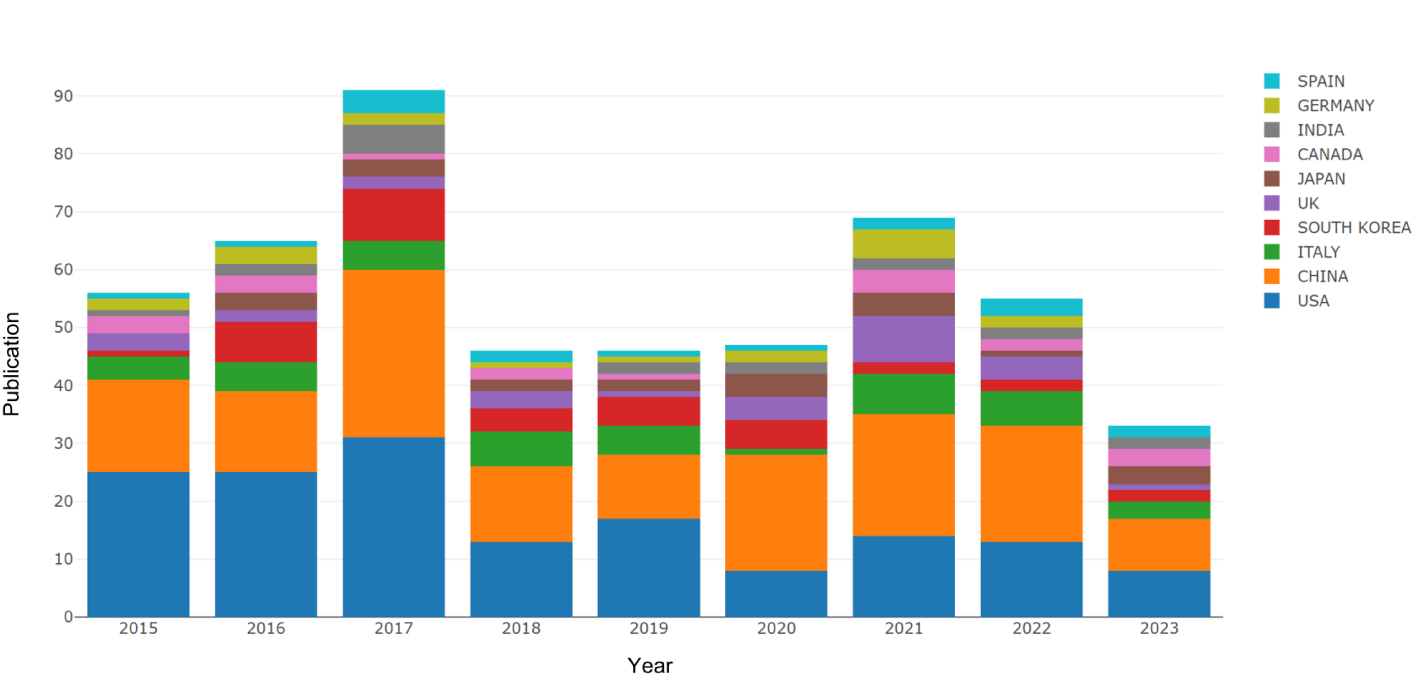
Annual Publication and the Contribution Characteristics of Each Country.

### 3.2. Countries analysis

In this study, a total of 77 countries participated in research related to mitochondria and prostate cancer. The annual publication trends of the top 10 countries between 2015 and 2023 are shown in Figure 1. The United States led in the number of publications until 2020, when China surpassed it. Detailed information about the top 10 countries is provided in Table 1. The United States ranked first in research productivity, publishing 154 papers with a total of 6,677 citations, and an average citation count of 43.36 per paper. China followed closely behind, with 138 papers published, accumulating 3,741 citations, and an average of 27.11 citations per paper. Italy ranked third, with 42 papers published, 2,509 citations, and an average of 59.74 citations per paper.

**Table 1.**
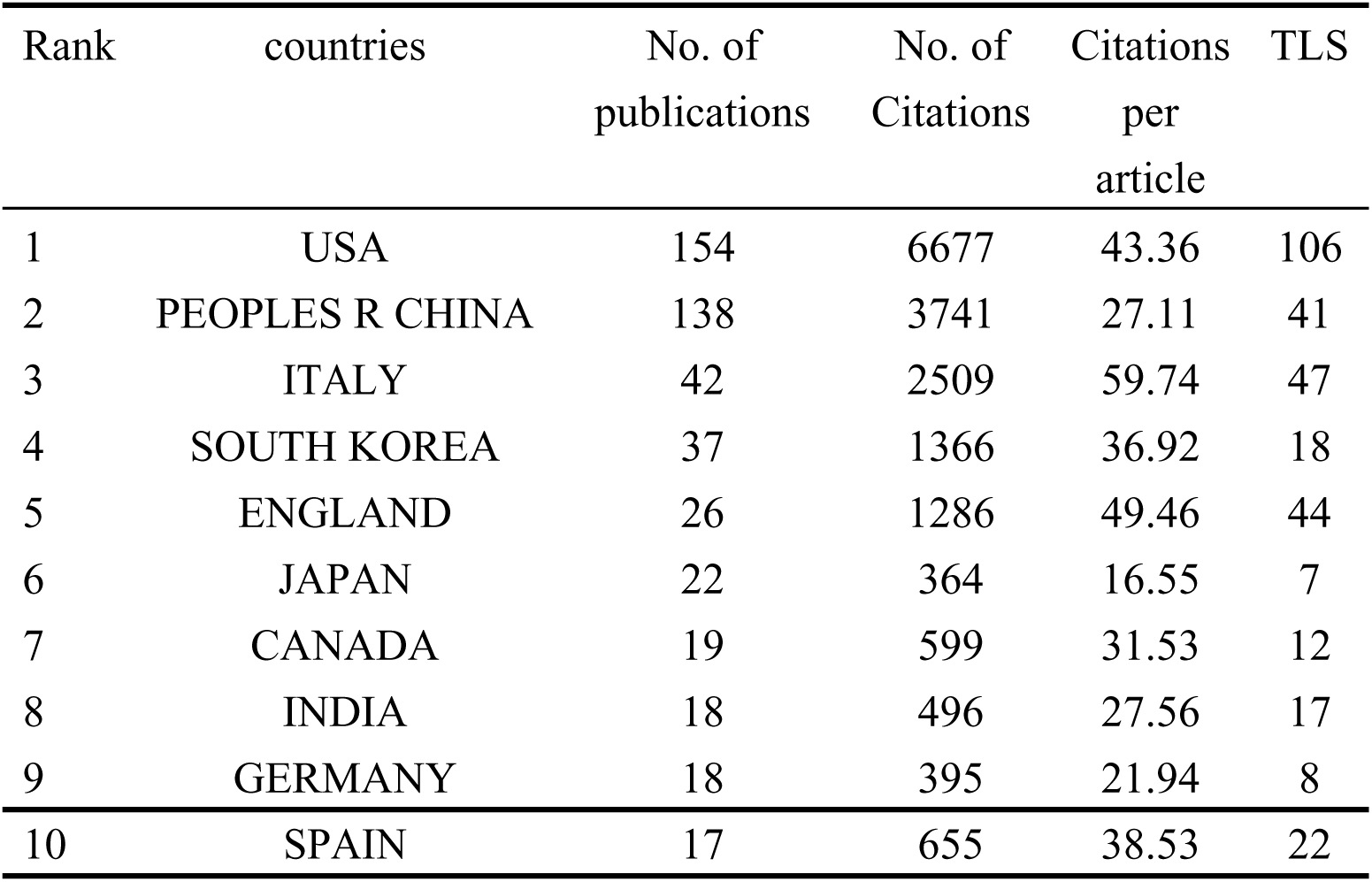
The top 10 productive countries.

The collaborative relationships between countries are depicted in Figure 2, with thicker lines indicating stronger collaborative ties. The United States has the most frequent collaborations with other countries/regions, followed by the United Kingdom. Notably, the collaboration between the United States and China is particularly close. The global collaboration network involving 26 countries, analyzed using VOSviewer, is shown in Figure 3. A minimum publication threshold of 5 was set, and TLS (Total Link Strength) represents the thickness of the connecting lines between nodes, reflecting the strength of international collaboration. The top four countries with the highest TLS values are the United States (TLS = 111), Italy (TLS = 49), the United Kingdom (TLS = 44), and China (TLS = 43).

**Figure 2.**
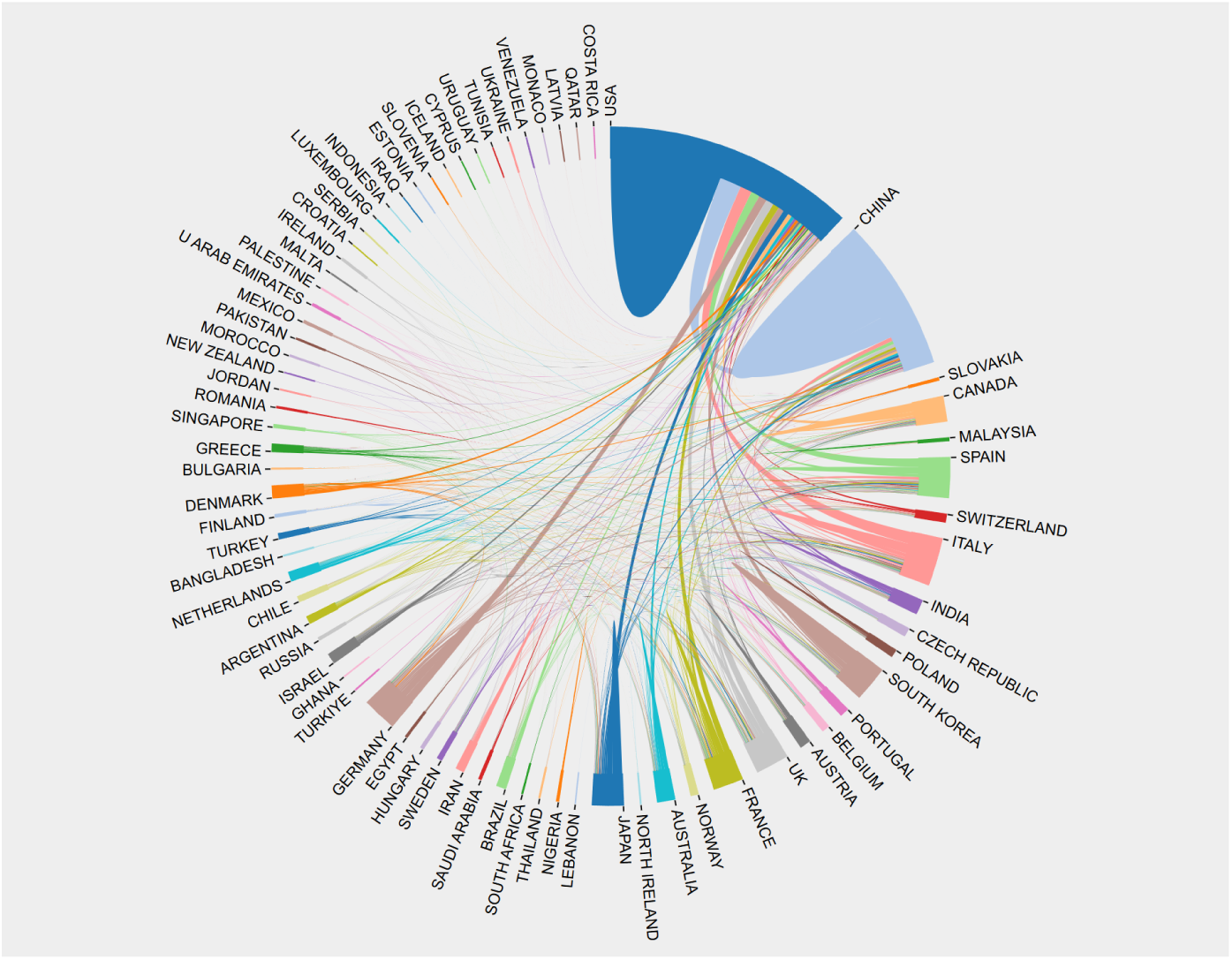
Global collaborative networks among countries.

**Figure 3.**
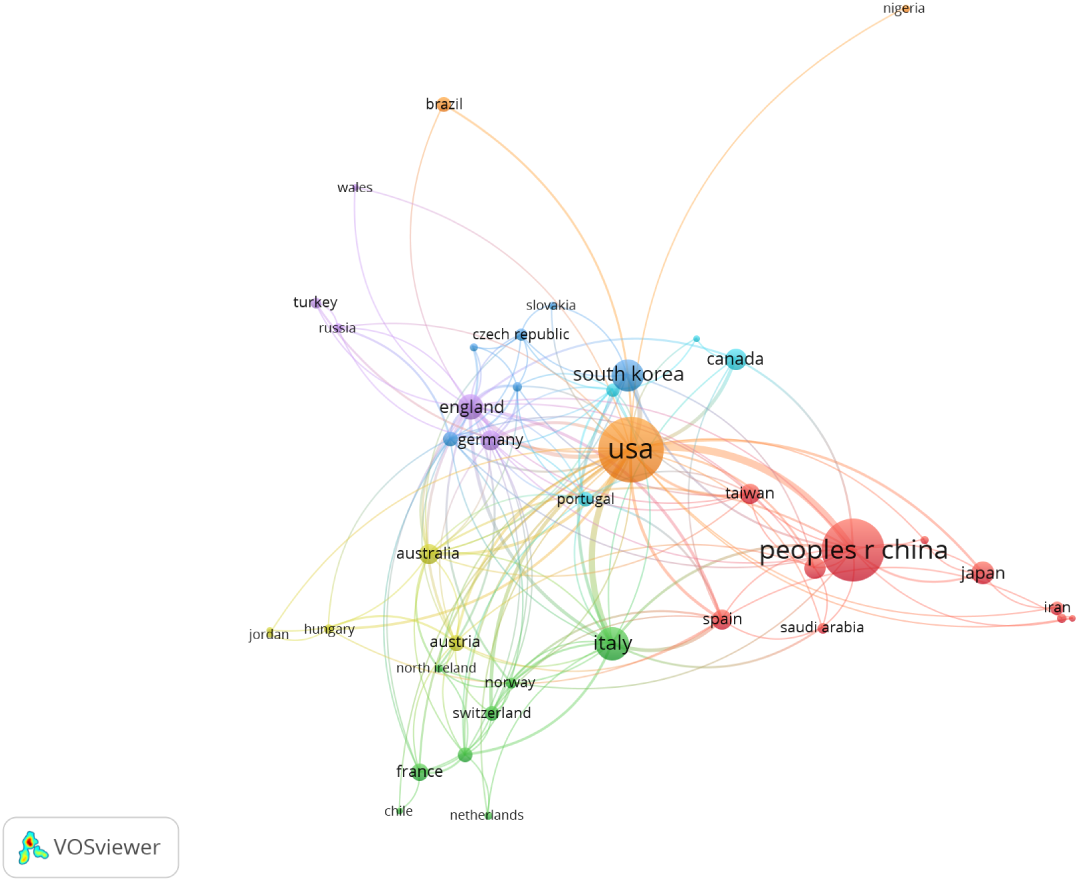
A citation network among various countries.

### 3.3. Contributions of top Institutions

During the study period, a total of 880 institutions participated in research related to mitochondria and prostate cancer. The top 10 institutions ranked by the number of publications are listed in Table 2. The Wistar Institute Anat & Biol was the largest contributor, publishing 24 articles (5.31% of the total). It was followed by Thomas Jefferson University and the University of Milan, which published 15 (3.32%) and 13 (2.88%) articles, respectively. In terms of citations, the Wistar Institute Anat & Biol ranked first with 1,301 citations, averaging 54.21 citations per paper. Thomas Jefferson University ranked second with 1,094 citations, averaging 72.93 citations per paper. Among the top 10 leading institutions, 4 are from the United States, 4 are from China, and 2 are from Italy.

**Table 2.**
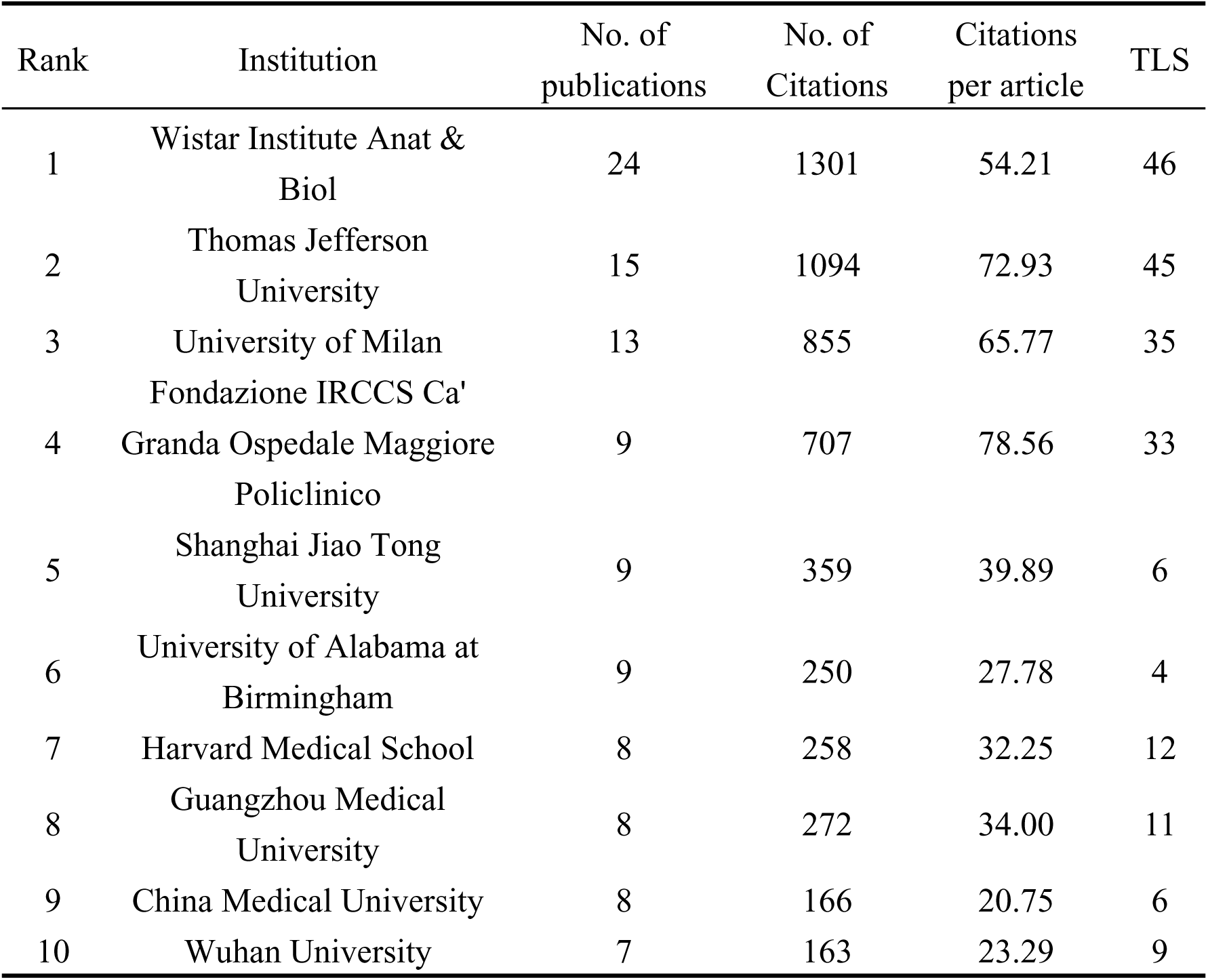
The top 10 productive institutes.

| Rank | Institution | No. of publications | No. of Citations | Citations per article | TLS |
| --- | --- | --- | --- | --- | --- |
| 1 | Wistar Institute Anat & Biol | 24 | 1301 | 54.21 | 46 |
| 2 | Thomas Jefferson University | 15 | 1094 | 72.93 | 45 |
| 3 | University of Milan | 13 | 855 | 65.77 | 35 |
| 4 | Fondazione IRCCS Ca' Granda Ospedale Maggiore Policlinico | 9 | 707 | 78.56 | 33 |
| 5 | Shanghai Jiao Tong University | 9 | 359 | 39.89 | 6 |
| 6 | University of Alabama at Birmingham | 9 | 250 | 27.78 | 4 |
| 7 | Harvard Medical School | 8 | 258 | 32.25 | 12 |
| 8 | Guangzhou Medical University | 8 | 272 | 34.00 | 11 |
| 9 | China Medical University | 8 | 166 | 20.75 | 6 |
| 10 | Wuhan University | 7 | 163 | 23.29 | 9 |

The institutional collaboration network map, generated by VOSviewer, is shown in Figure 4. It displays institutions with more than three publications in the field, excluding institutions without any collaboration. The connections between institutions represent the strength of their collaboration. This network includes 71 institutions, divided into 9 groups based on collaboration strength. In terms of size and TLS, the Wistar Institute Anat & Biol is the largest node, having published 24 articles with a TLS value of 46, indicating the institution’s greatest influence in international collaboration.

**Figure 4.**
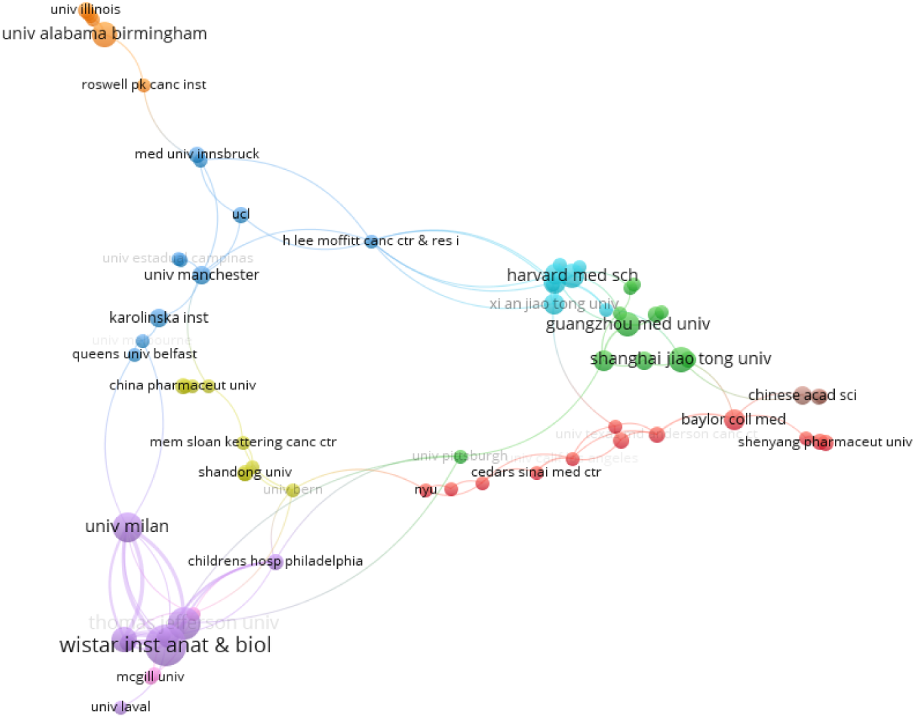
Network map of co-institutions.

### 3.4. Contributions of top Journals

A total of 241 journals published the 452 papers, with 16 journals publishing at least 5 papers each. In Table 3, the top 10 journals with the highest publication volume collectively published 99 papers, accounting for 21.90% of the total publications. The three journals with the highest publication volumes were Oncotarget (25 papers, 5.53%), International Journal of Molecular Sciences (14 papers, 3.10%), and Frontiers in Oncology (12 papers, 2.65%). Among these, Oncotarget had the highest total citation count, with 1,101 citations, significantly higher than those of other journals. However, Cell Death & Disease had the highest average citation count per paper, with an average of 103.29 citations, ranking first among all journals. This suggests that the papers published in Cell Death & Disease are of particularly high quality, indicating their significant impact in the field.

**Table 3.**
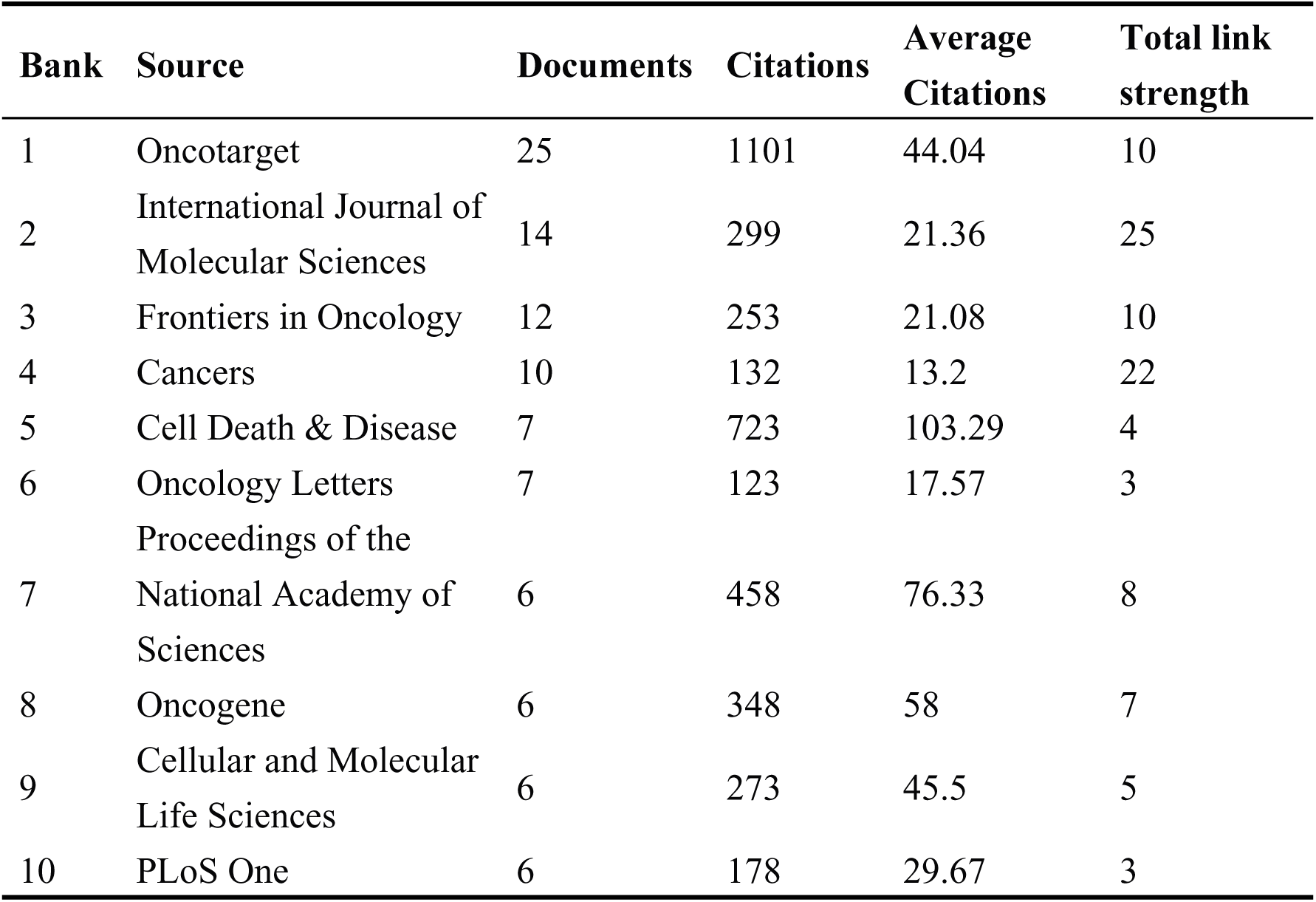
The top 10 productive Journals.

### 3.5. Analysis of Authors and Co-Cited Authors

Our analysis identified 3,179 primary researchers and 20,657 co-cited authors. The top 10 authors who have published extensively and have been co-cited in the field of mitochondria and prostate cancer are listed in Table 4. These top 10 high-productivity authors collectively published 119 papers, accounting for 26.33% of the total publications. Dario C. Altieri ranks first with 24 publications and 1,301 citations, followed by M. Cecilia Caino (13 publications, 938 citations), Lucia R. Languino (13 publications, 903 citations), and Jae Ho Seo (13 publications, 808 citations).

**Table 4.**
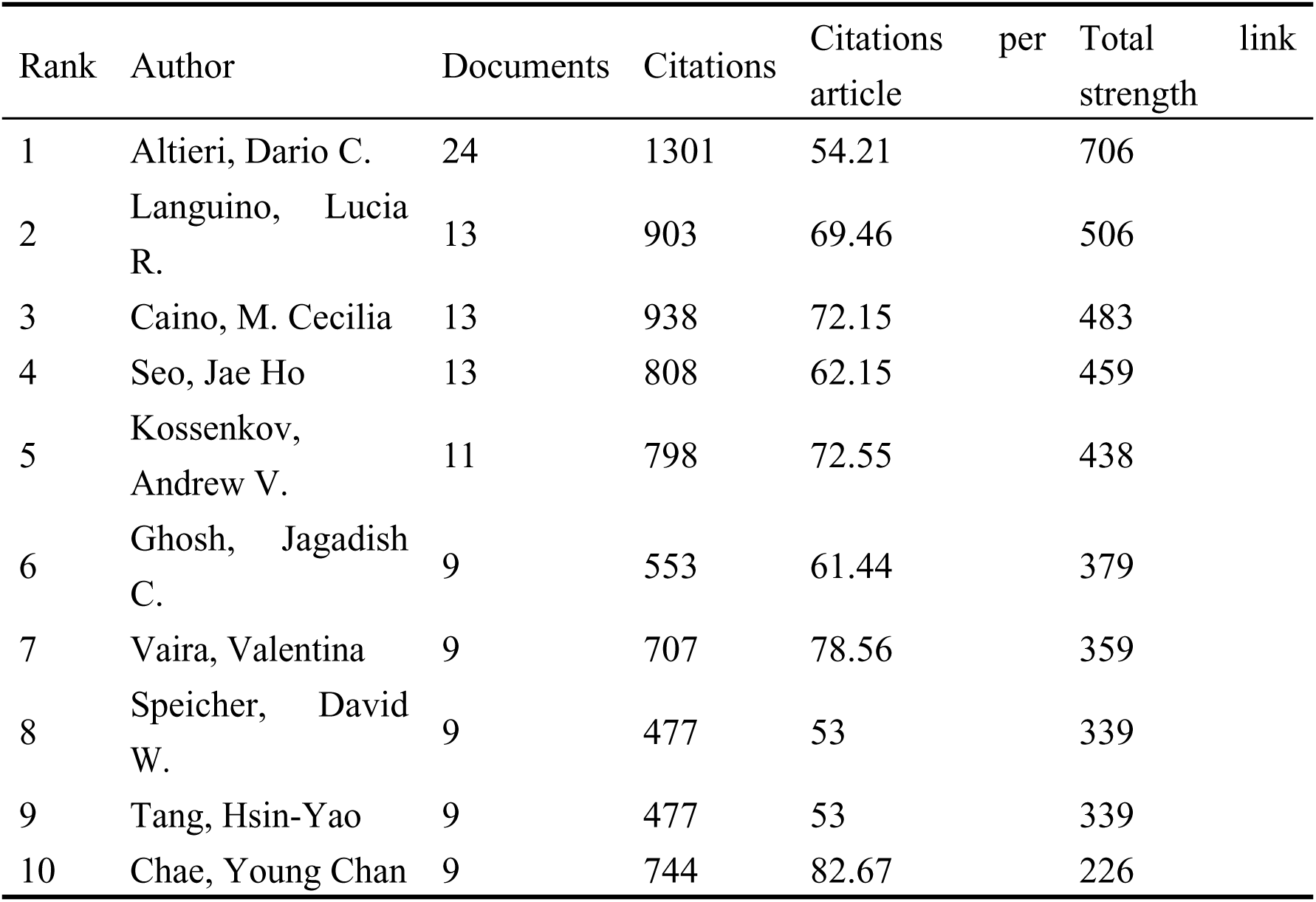
The top 10 productive authors.

In Figure 5A, the collaboration visualization network generated by VOSviewer is presented, illustrating the collaboration relationships of authors who have published more than three papers in the field. Authors without collaborations are excluded from the visualization. Each node represents an author, with the size of the node indicating the number of published papers, and the links between nodes representing collaboration relationships. The network includes 22 authors, each having published more than three papers, forming four author clusters. Among these authors, Dario C. Altieri has the most collaborators, mainly working with U.S.-based researchers, while Chinese researchers have more limited collaboration in this field.

**Figure 5.**
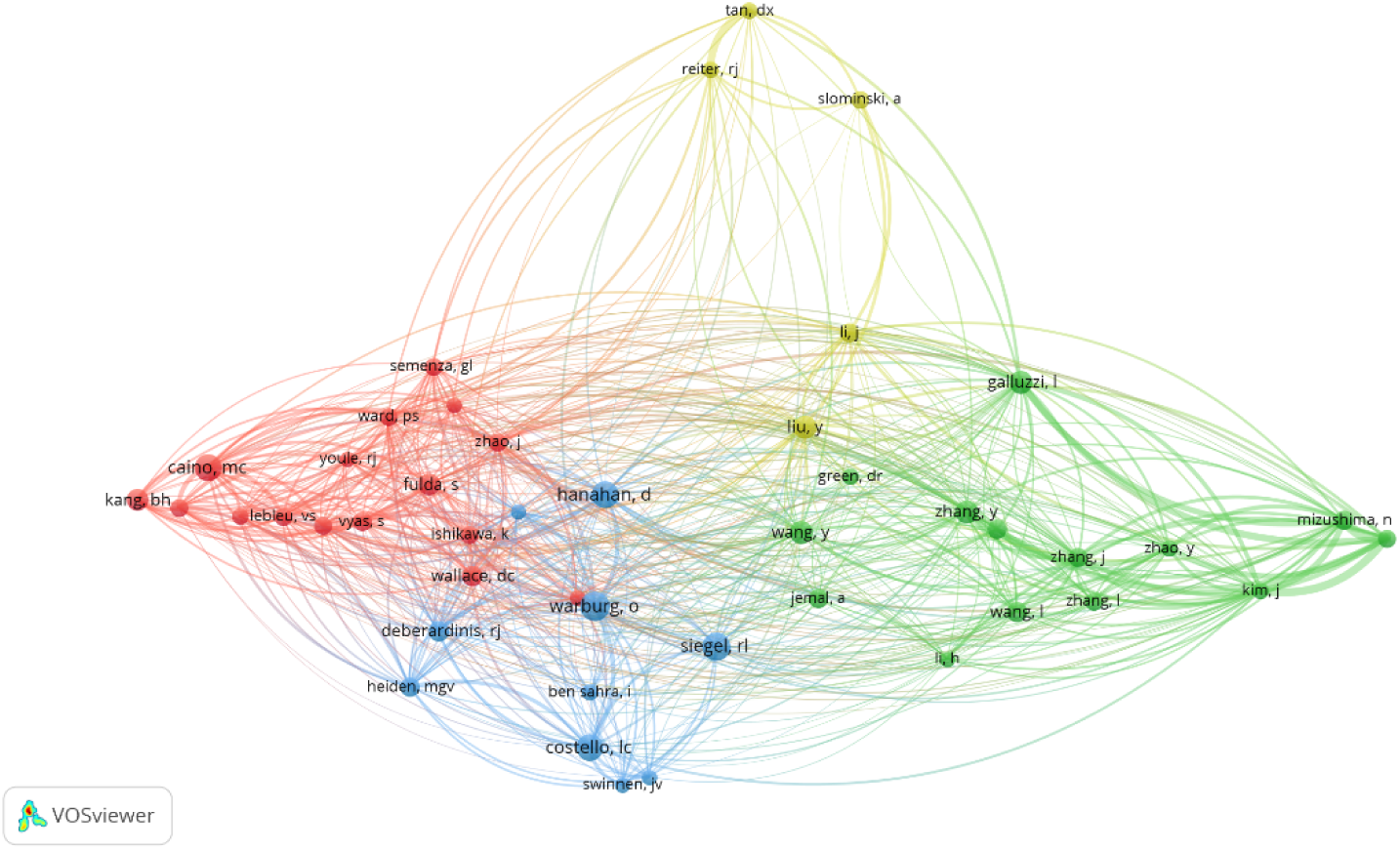
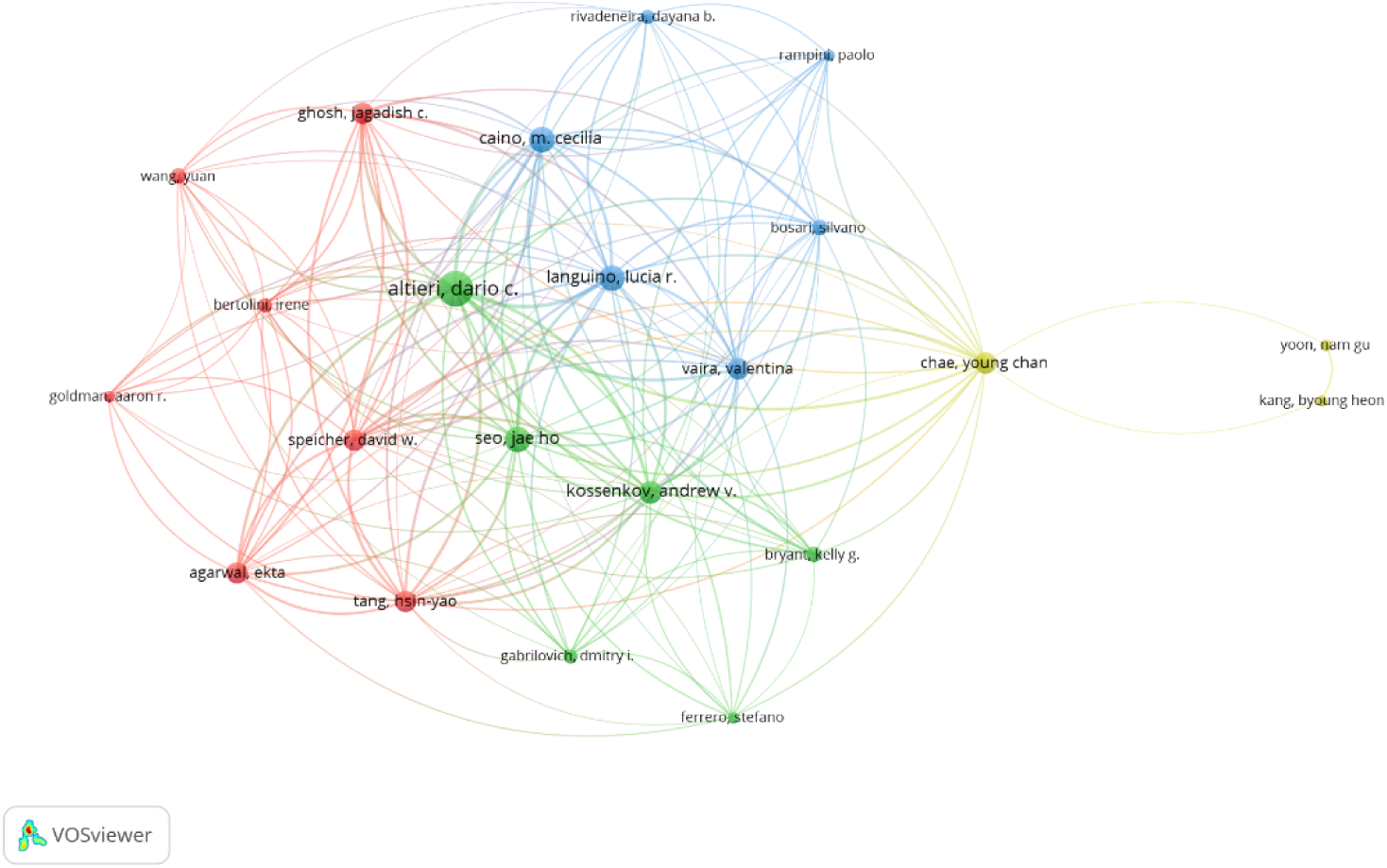
Network maps of co-authorship(A) and co-citation(B) authors analyses.

Further analysis of co-citation patterns is shown in Figure 5B, which presents a network of 72 authors who have been cited more than 15 times. The node size reflects the number of citations each author has received, and the links represent co-citation relationships between authors. TLS indicates an author’s influence on peers in the field. Among the most co-cited authors, O. Warburg ranks first with 72 citations (TLS = 595), followed by I. C. Costello (61 citations, TLS = 526) and M. C. Caino (58 citations, TLS = 543).

### 3.6. Analysis of references

According to Table 5, the top 10 most cited articles have been cited a total of 6,984 times, accounting for 39.29% of the total citations. Among these, the two articles by Klionsky, Daniel J. — “Guidelines for the Use and Interpretation of Assays for Monitoring Autophagy (3rd edition)” published in 2016 and “Guidelines for the Use and Interpretation of Assays for Monitoring Autophagy (4th edition)” published in 2020 — have received 3,703 and 936 citations, respectively. The average annual citation counts for these two articles are 411.44 and 234, respectively, significantly higher than those of other articles, highlighting their substantial influence within the field.

**Table 5.**
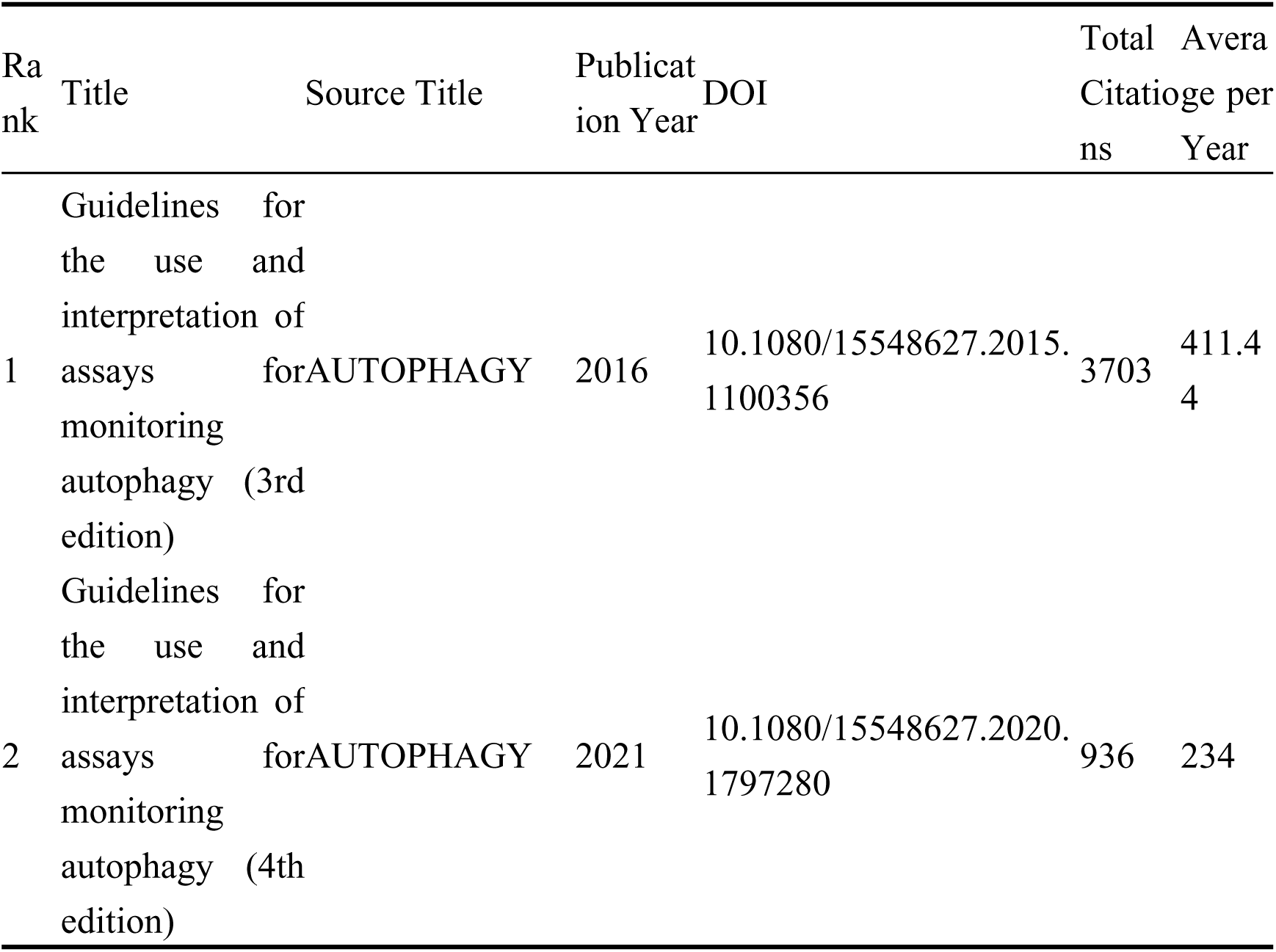
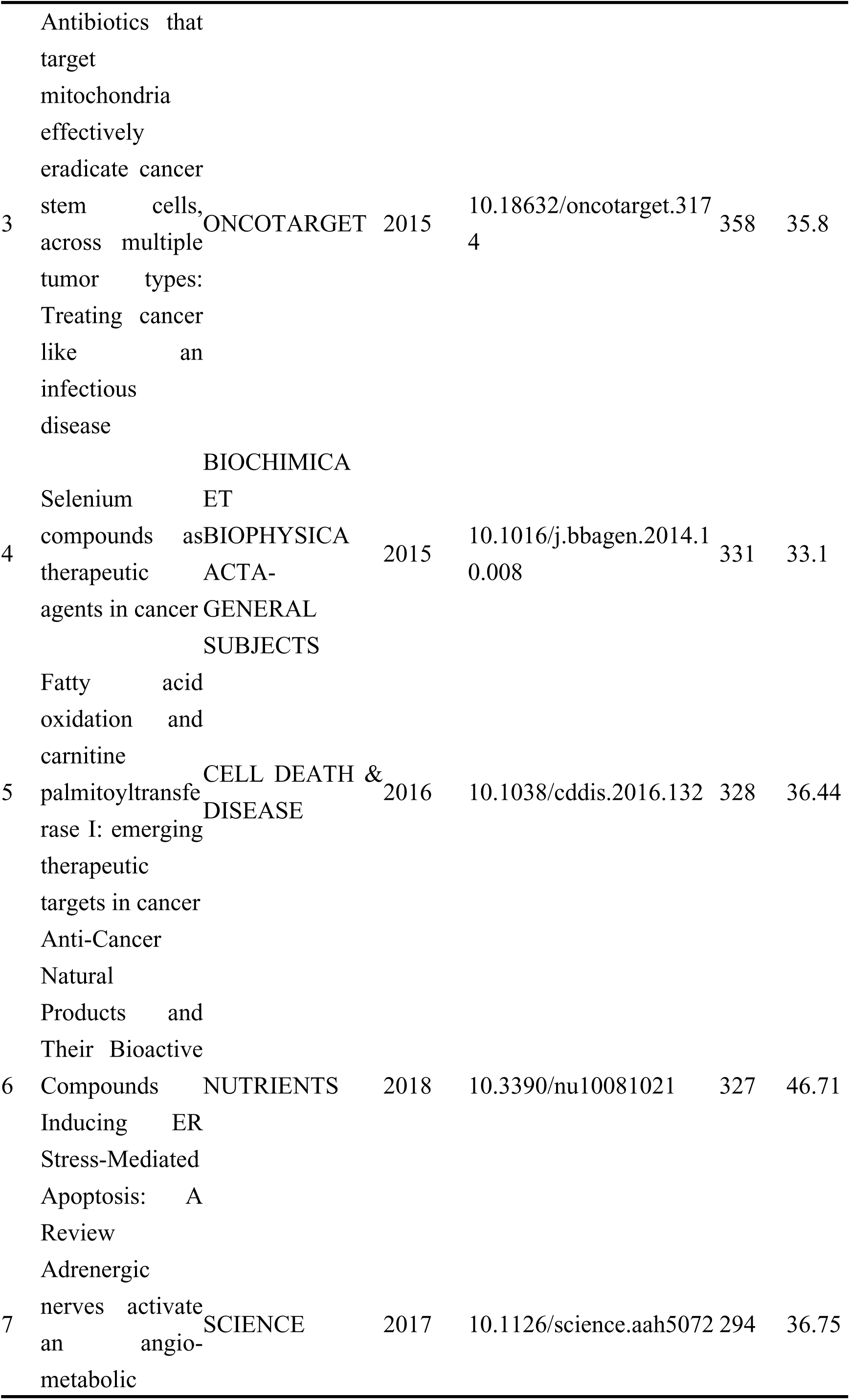
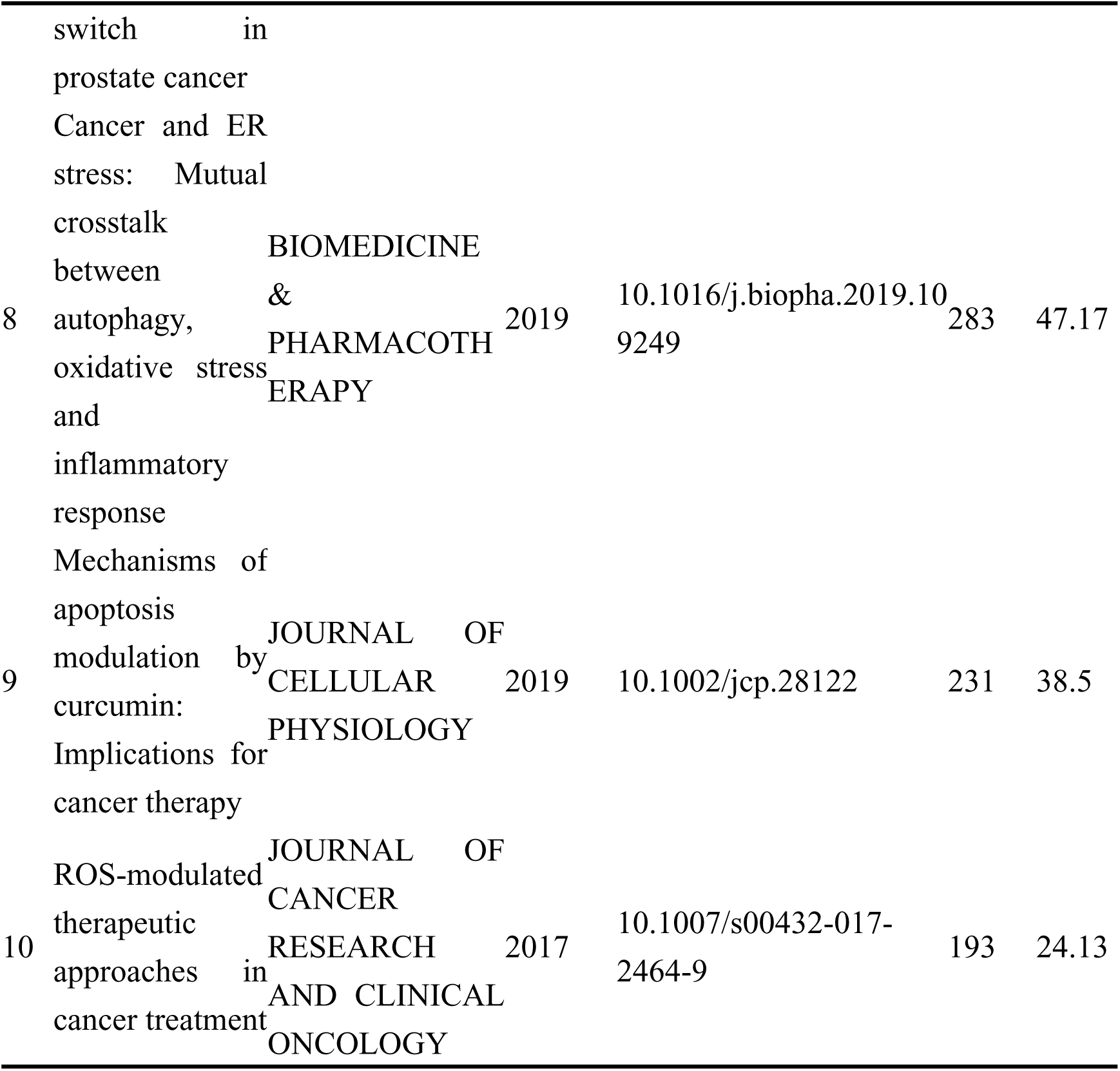
The top 10 most cited references.

Using CiteSpace software, we generated a co-citation network map of references and conducted a clustering analysis, which revealed 16 distinct clusters (Figure 6A). The modularity Q value was 0.7309, and the mean silhouette value was 0.8869, both exceeding 0.5, indicating a high degree of validity in the clustering results. Cluster 1 was labeled “metastasis” (#0), and Cluster 3 was labeled “metabolomics” (#4), suggesting that metastasis and metabolism are key research areas regarding the role of mitochondria in prostate cancer. Cluster 6 was labeled “autophagic degradation” (#7), indicating that autophagic degradation of mitochondria is also a significant current research focus. Cluster 10 was labeled “therapy resistance” (#9), underscoring the ongoing challenge of drug resistance in prostate cancer research.

**Figure 6.**
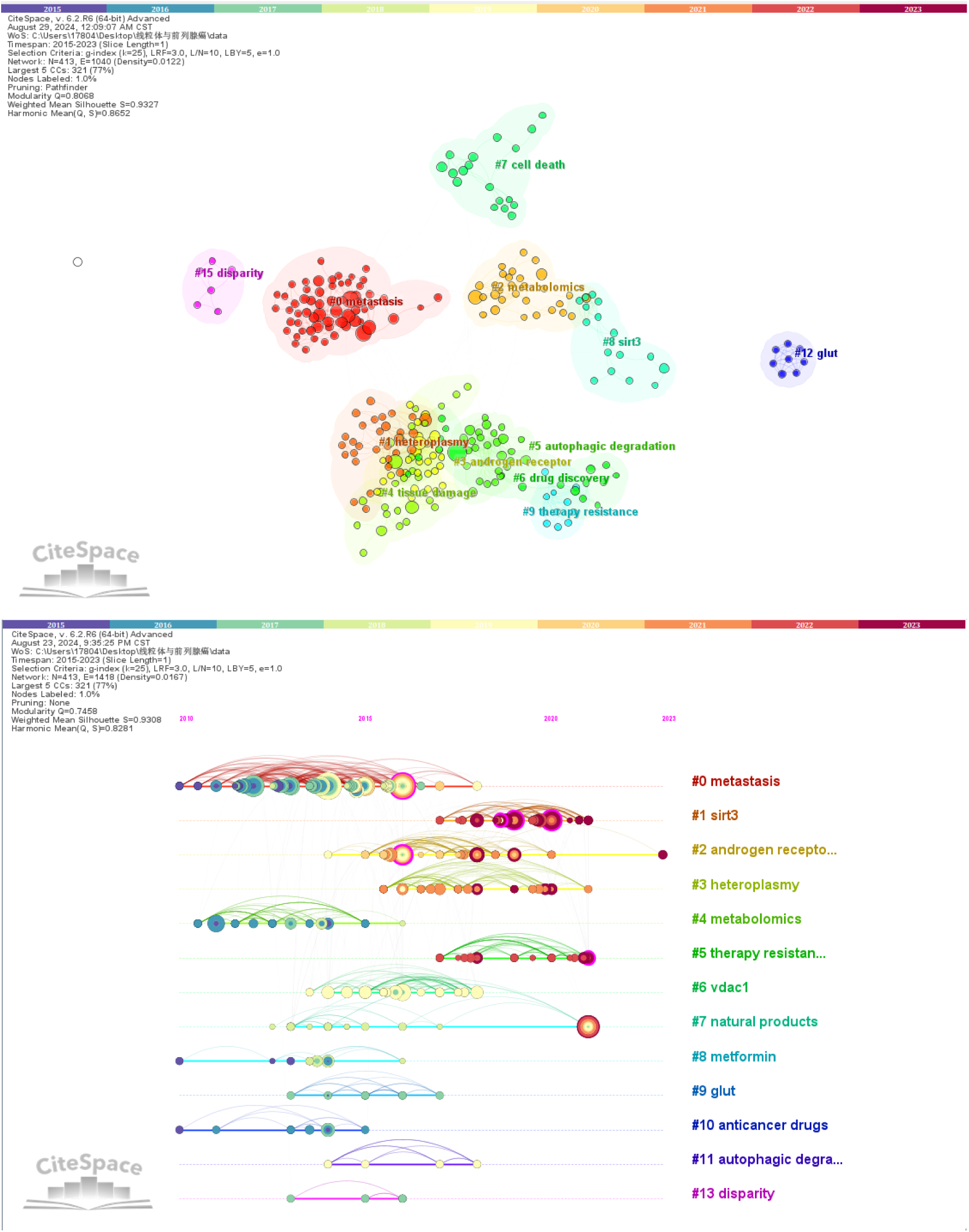
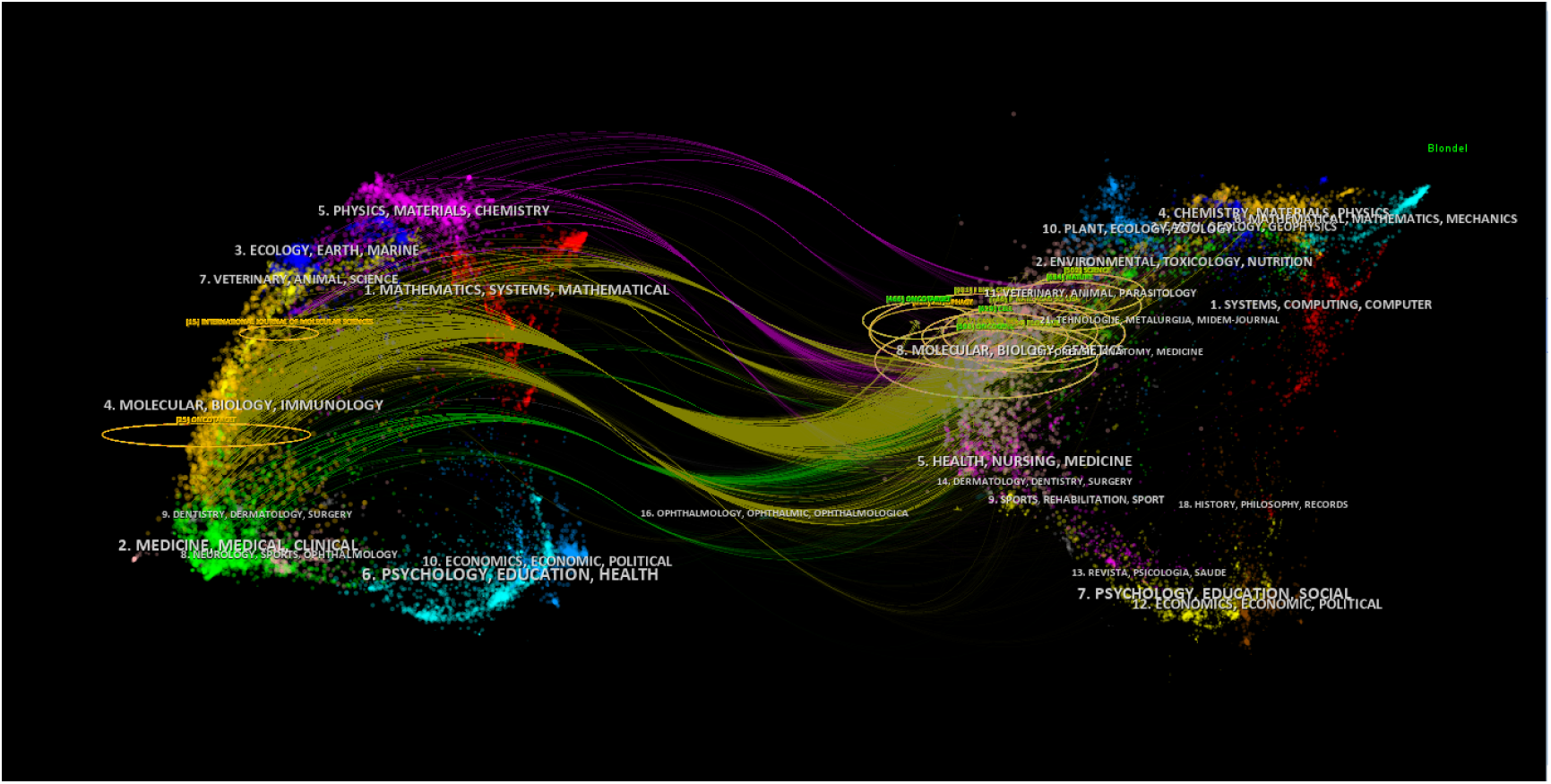
Network maps of Cluster view (A) and timeline view (B) and dual-map overlay. (C)of co-citation references.

Furthermore, we constructed a timeline visualization map of co-cited references to provide insights into the evolution of research topics in this field (Figure 6B). In this map, nodes on the same line are distinguished by different colors to represent articles from different years; nodes on the left represent earlier references, while those on the right represent more recent publications. Our analysis reveals that early research predominantly focused on “metastasis,” “metabolomics,” and “anticancer drugs,” while contemporary research trends are primarily concentrated on “therapy resistance,” “androgen receptor,” “heteroplasmy,” and “natural products.”

Finally, we created a dual-map overlay of the literature to visualize the distribution of topics within each article (Figure 6C). The map superimposes the dual graphs of citing and cited journals, revealing three primary citation paths represented by yellow, green, and pink lines, which illustrate the flow of knowledge. Specifically, citing journals are mainly concentrated in the fields of molecular biology, biology, immunology, medicine, and clinical research, while cited journals are predominantly found in the fields of molecular biology, biomedical sciences, health sciences, and nursing.

### 3.7. Analysis of keywords

It is noteworthy that keywords reflect the central themes and research directions of publications. Therefore, we utilized VOSviewer to construct a keyword network map. As shown in Figure 7, these keywords are divided into nine clusters (red, green, blue, yellow, purple, cyan, brown, pink, and tan). Specifically, Cluster 1 (green) contains 13 items, covering topics such as mitochondria, SIRT3, and oxidative phosphorylation; Cluster 2 (red) includes 12 items, focusing on research directions like doxorubicin and combination therapy; Cluster 3 (blue) comprises 10 items, such as cell cycle, melatonin, and cytotoxicity; Cluster 4 (yellow) contains 9 items, primarily related to DU145 (a prostate cancer cell line), castration-resistant prostate cancer, and the androgen receptor; Cluster 5 (purple) consists of 8 items, addressing research topics such as mitochondrial DNA, oxidative stress, and bioenergetics; Cluster 6 (pink) includes 8 items, revolving around themes such as autophagy, the Warburg effect, and hypoxia; Cluster 7 (cyan) consists of 7 items, with keywords including PI3K, HSP90, and metformin; Cluster 8 (brown) also contains 7 items, with a focus on reactive oxygen species, MAPK, and antioxidants; and finally, Cluster 9 (tan) consists of 6 items, related to research on P53 and proliferation.

**Figure 7.**
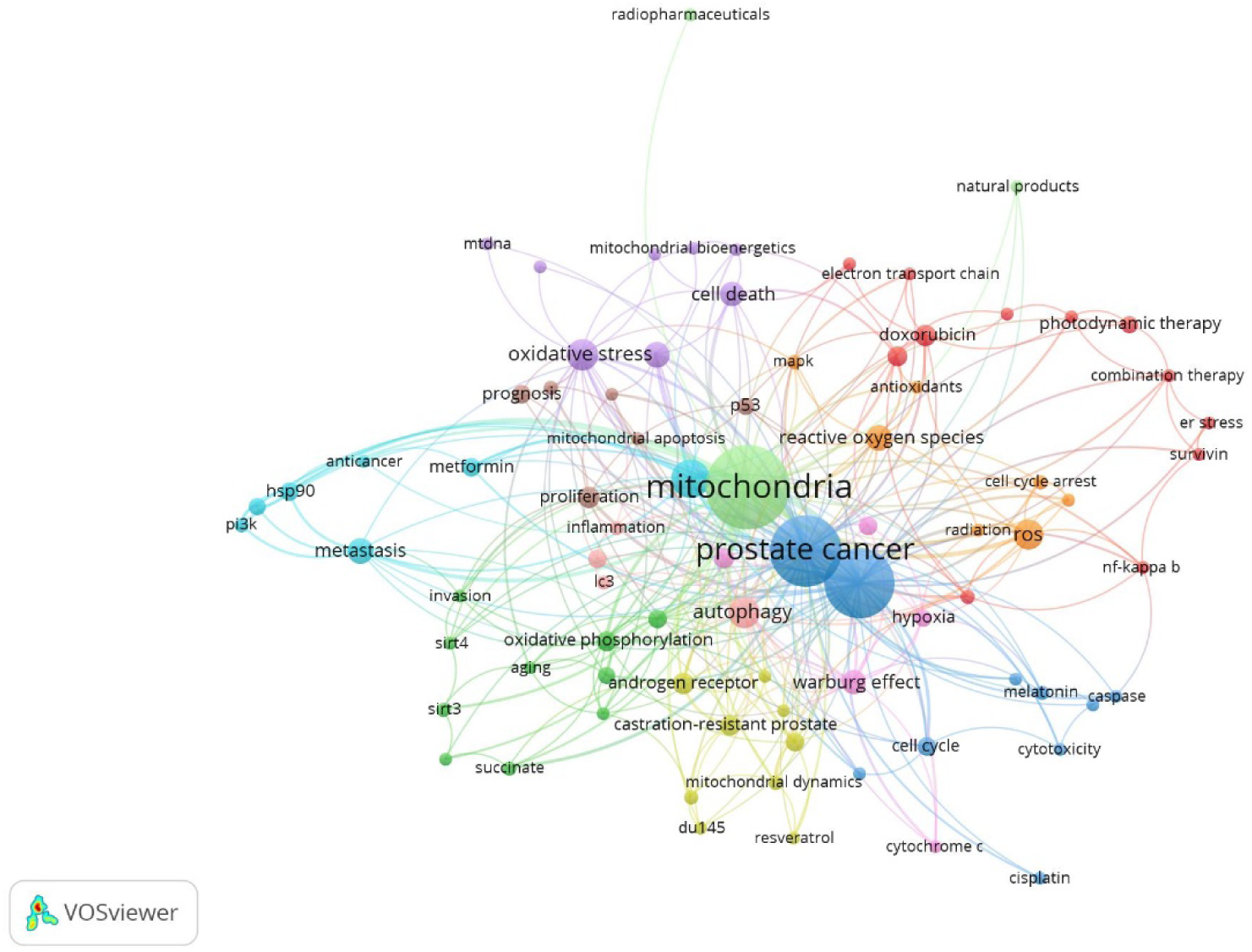
Network map of co-occurrence keyword.

Additionally, we created a timeline visualization map of co-cited keywords to better understand the evolution of research topics. In Figure 8, nodes on the same timeline are distinguished by different colors to represent different years. Nodes on the far left correspond to earlier hot-topic keywords, while those on the right represent more recent research trends. Our analysis indicates that early research keywords primarily focused on “inflammation,” “growth,” and “reactive oxygen species, “ while current research hotspots have shifted towards “drug resistance,” “radiotherapy resistance,” and “therapy.”

**Figure 8.**
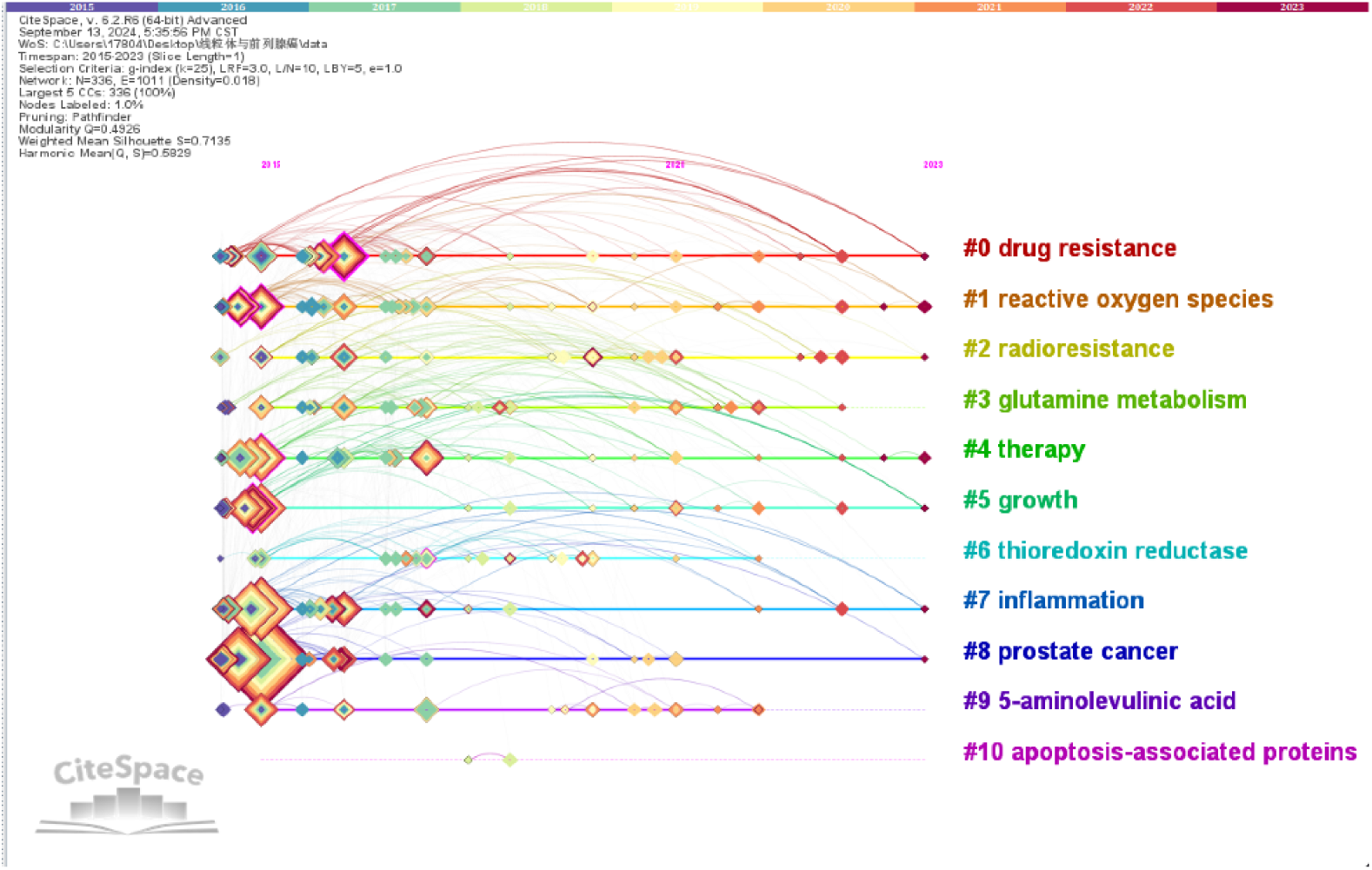
Keyword timezone map.

### 3.8. Summarization of experiments information

We reviewed 241 experimental studies, of which 110 were in vivo experiments and 131 were in vitro or ex vivo studies (Figure 9). Among the studies on mitochondrial-targeted therapy for prostate cancer, 152 articles were identified, with 29 focusing on targeted therapy and combination therapy (Figure 10). By analyzing genes closely related to mitochondrial functions and prostate cancer progression, we found that LONP1 (Lon Peptidase 1, Mitochondrial), Parkin, and DRP1(Dynamin-related protein 1) were the three most frequently mentioned genes.

**Figure 9.**
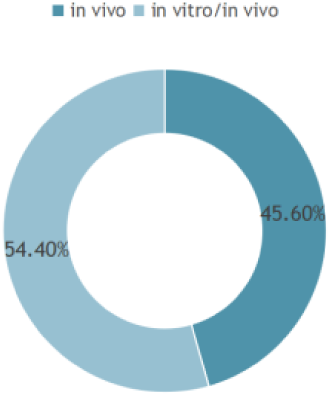
Statistics of experimental methods.

**Figure 10.**
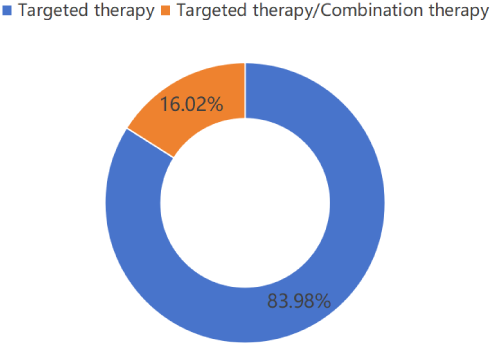
Statistics of therapy methods.

Parkin is an E3 ubiquitin ligase that regulates mitochondrial function and metabolism through multiple mechanisms, thereby inhibiting prostate cancer development. These mechanisms include regulating mitochondrial dynamics, influencing mitochondrial protein function, and modulating related signaling pathways through ubiquitination modification[9]. DRP1 is a GTPase associated with dynamin, primarily involved in regulating mitochondrial morphology, particularly playing a key role in mitochondrial fission within the cell, which significantly impacts the progression of prostate cancer[10].

LONP1 is an ATP-dependent protease located in the mitochondrial matrix. LONP1 functions through various mechanisms, including maintaining mitochondrial function, regulating cell metabolism, inhibiting apoptosis, modulating oxidative stress, and promoting tumor metastasis, thereby playing a crucial role in cancer initiation and progression[11–13]. However, research on the relationship between LONP1 and prostate cancer remains limited. To investigate the correlation and underlying mechanisms of LONP1 in prostate cancer, we conducted relevant studies.

Figure 11 presents the expression levels of the LONP1 gene in prostate adenocarcinoma (PRAD) from the TCGA database, comparing the expression levels based on sample type. The data show that the expression of the LONP1 gene is upregulated in prostate adenocarcinoma and may be associated with tumor initiation and progression in prostate adenocarcinoma. Furthermore, statistical tests revealed that the difference between the two groups is statistically significant (p < 0.001).

**Figure 11.**
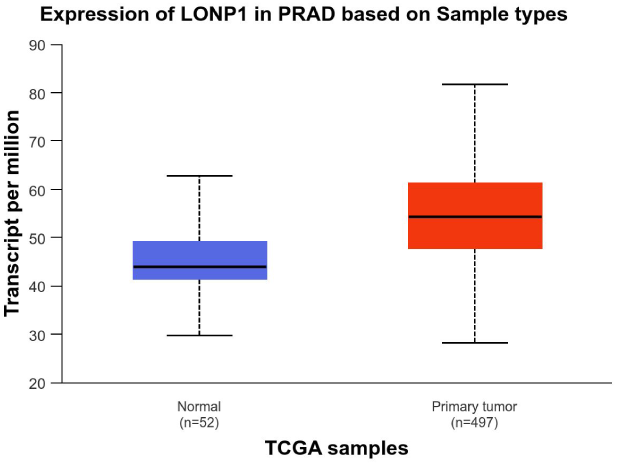
Expression of LONP1 in PRAD and normal tissues based on TCGA.

In Figure 12, the expression of the LONP1 gene across various cancer types in the TCGA (The Cancer Genome Atlas) database is further analyzed. The expression level of LONP1 in PRAD (prostate adenocarcinoma) is moderate relative to other tumor types.

**Figure 12.**
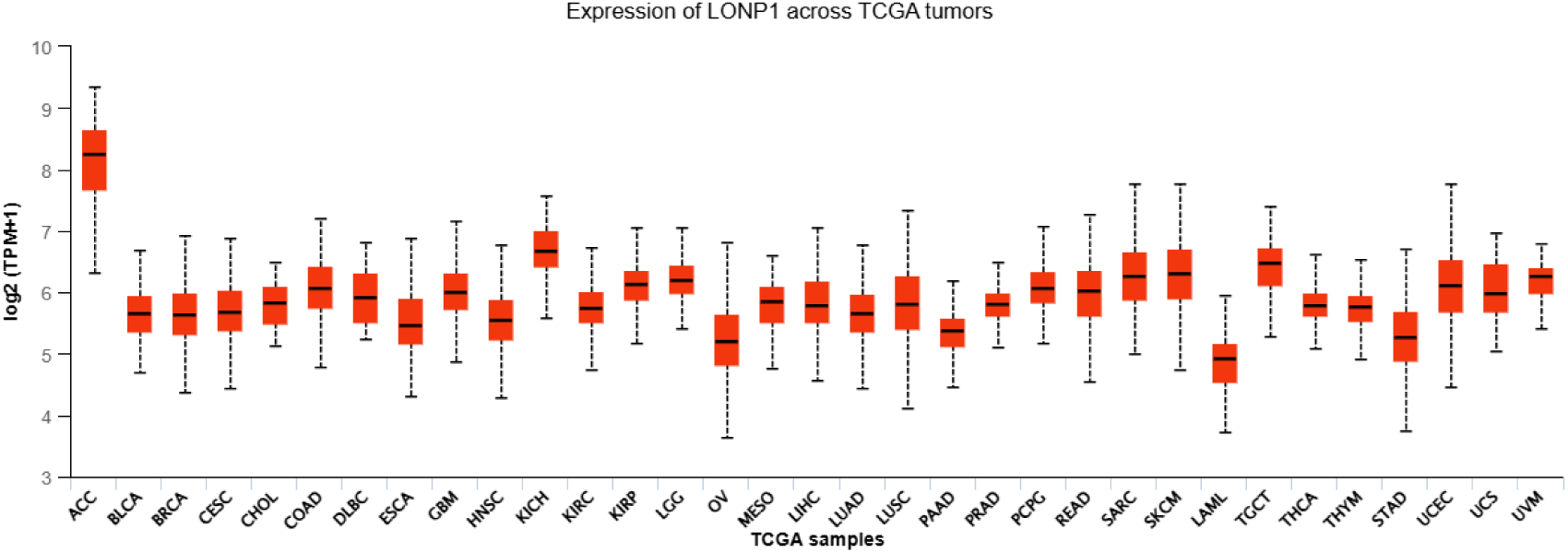
Expression of LONP1 across TCGA tumor.

The biological processes that the LONP1 gene is involved in are illustrated in Figure 13, based on Gene Ontology (GO) analysis. These processes include mitochondrial DNA metabolism, maintenance of the mitochondrial genome, chaperone-mediated protein complex assembly, protein quality control, and the degradation of cellular proteins. This information contributes to a deeper understanding of the potential intracellular functions of LONP1.

**Figure 13.**
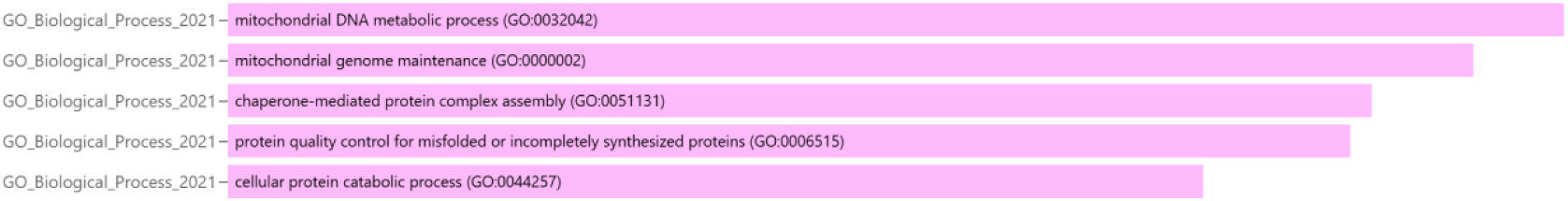
Biological processes associated with the LONP1 gene.

Figure 14 presents the Spearman correlation data between the LONP1 gene and various cancer-related pathways in prostate cancer (PRAD). These pathways include apoptosis, cell cycle, DNA damage, epithelial-mesenchymal transition (EMT), androgen receptor (AR), estrogen receptor (ER), PI3K/AKT, RAS/MAPK, receptor tyrosine kinase (RTK), and TSC/mTOR.

**Figure 14.**
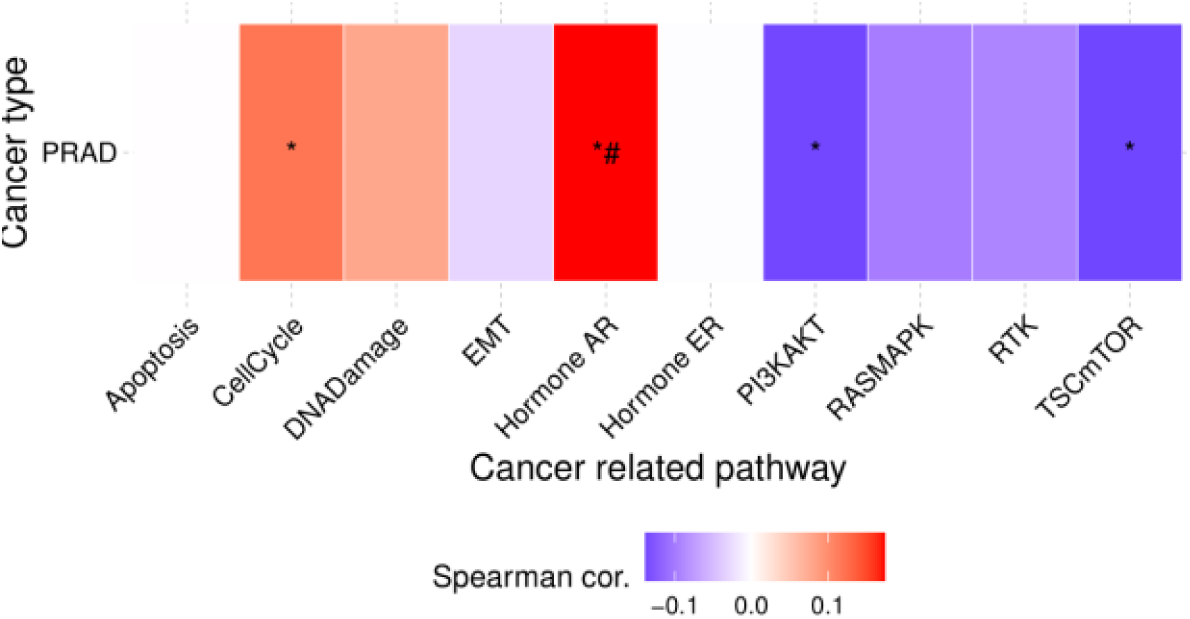
Spearman correlation of LONP1 gene with prostate cancer-related pathways.

Additionally, LONP1 is involved in regulating the expression of mitochondrial genes, affecting the replication and transcription of mitochondrial DNA (mtDNA) as well as the stability of related regulatory factors, thereby controlling the synthesis of mitochondrial-encoded proteins. Based on these functions, LONP1 has the potential to serve as a promising biomarker and therapeutic target for prostate cancer[14]. We also investigated the relationship between LONP1 and anticancer drug sensitivity to further explore its potential applications in therapy.

Figures 15A and 15B show the correlation between LONP1 gene expression and drug sensitivity from the GDSC (Genomics of Drug Sensitivity in Cancer) database. As indicated by the figures, the expression of LONP1 has a significant negative correlation with several drugs, including Vincristine, GSK461364 (Parthenate), Methotrexate, and THZ-2-102-1THZ2. LONP1 expression shows varying degrees of correlation with the sensitivity to these drugs, providing clues for future research on drug targeting.

**Figure 15.**
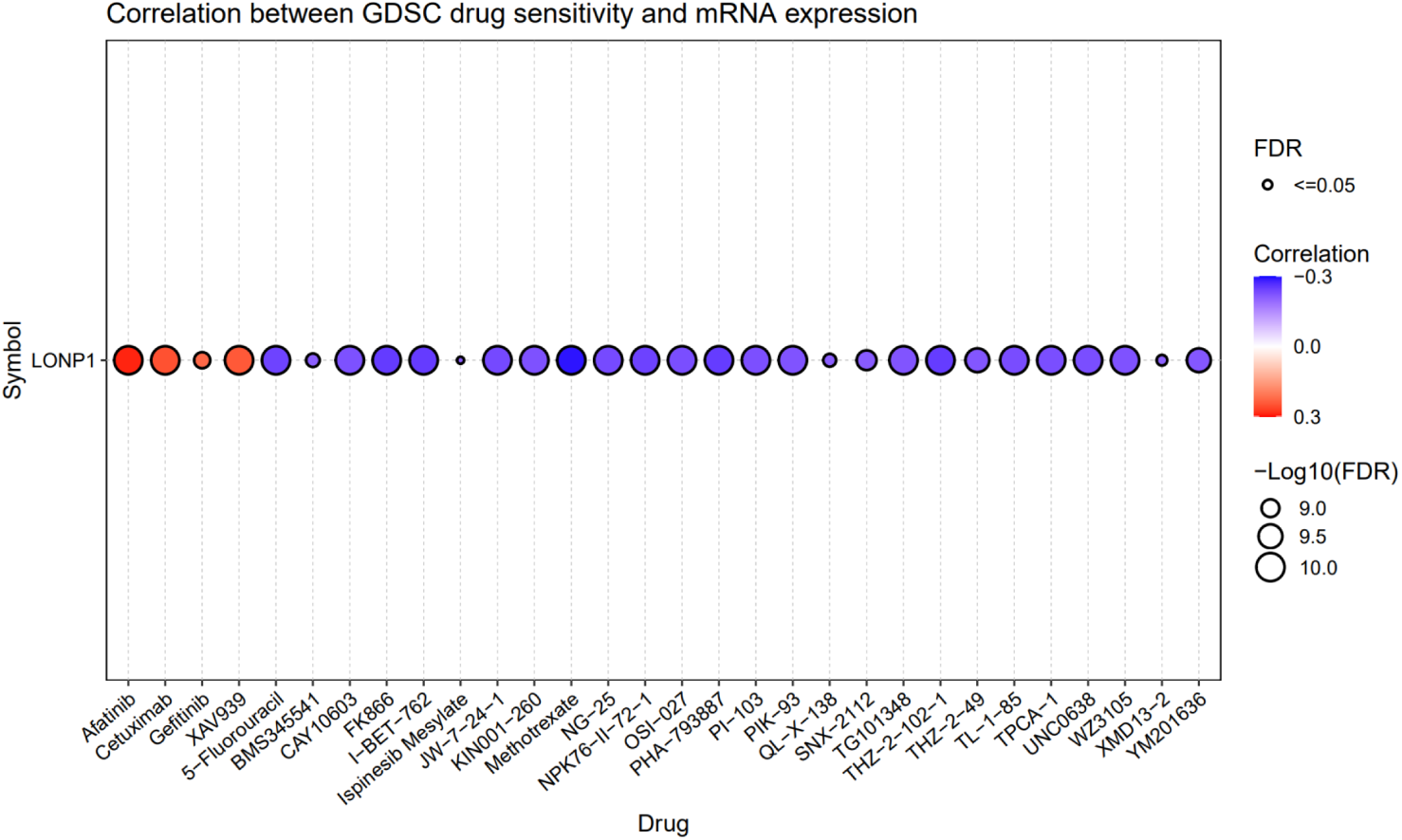
A and B Correlation analysis between GDSC drug sensitivity and LONP1 mRNA expression.

However, there are some limitations to the use of these small-molecule drugs in treatment. Vincristine, GSK461364, Methotrexate, and THZ-2-102-1THZ2 are all chemotherapy drugs. First, patients may develop resistance to these treatments, leading to a gradual reduction in drug efficacy[15]. Additionally, these treatments may cause various side effects, such as myelosuppression, gastrointestinal reactions, increased cardiovascular disease risk, hepatotoxicity, nephrotoxicity, and more[16]. Some drugs, such as Vincristine and Parthenate, can also induce peripheral neuropathy, which significantly affects patients’ quality of life[17]. Furthermore, due to individual differences, treatment plans must be personalized according to the patient’s specific condition, making treatment decisions more complex.

In cancer treatment, compared to traditional molecular drug therapies, targeted antibody therapies offer the following advantages. Antibody treatments are more specific and help reduce the common side effects associated with conventional drug therapies. Targeted antibody drugs can specifically recognize and bind to specific antigens on target cells or inside cells, avoiding cross-reactivity with other antigens, thus minimizing damage to normal cells[18]. For example, trastuzumab specifically recognizes and binds to the HER-2 antigen on the surface of breast cancer cells, exhibiting highly specific cytotoxic effects on HER-2 positive breast cancer cells, while having minimal impact on normal cells. Bevacizumab is used to treat a variety of solid tumors, such as colorectal cancer and non-small cell lung cancer. These drugs do not cause severe side effects like myelosuppression and hair loss commonly seen with chemotherapy. Although common side effects such as hypertension and proteinuria may occur, they are relatively mild and most patients can tolerate them[19,20].

### 3.9. Virtual design of specific antibodies

Current research on antibody therapeutics targeting LONP1 remains relatively scarce. To address this knowledge gap, we utilized the GeoBiologics platform to design and screen fully human antibodies against LONP1. The GeoBiologics platform integrates critical functionalities including antibody structure prediction, affinity assessment, and immunogenicity profiling, supporting de novo design and multi-objective optimization to deliver an AI-powered comprehensive workflow from target identification to preclinical candidate characterization. Our platform achieves comparable success rates to AlphaFold3 in complex structure prediction, demonstrating twice the accuracy of AlphaFold2 and significantly outperforming conventional computational tools such as MOE.

The generative AI-driven antibody optimization framework implemented in this platform enabled rapid precision engineering of the 8G3 antibody, achieving a remarkable 1000-1500-fold enhancement in neutralizing activity against the emerging JN.1 viral variant[21]. This breakthrough not only validates the transformative potential and broad applicability of AI-powered antibody engineering but also establishes an efficient alternative to conventional antibody discovery methodologies. Capitalizing on the advanced capabilities of GeoBiologics, we employed its Targeted CDR Library Design module to construct antigen-specific complementarity-determining region (CDR) libraries directed against LONP1.

During the design process, we precisely defined the antigen epitopes and antibody scaffold, successfully predicting a monoclonal antibody with high affinity and specificity. The selected antigen is LONP1, with its epitope located at the 421–558 amino acid residues of the C chain. The antibody scaffold is based on the well-known monoclonal antibody Trastuzumab (1n8z), which is commonly used to construct new antibody candidates. We used the GeoFlowDesign model to guide the antibody design and screened 100 different antibody designs. Each design underwent preliminary affinity and specificity evaluation in a simulated environment.

Through systematic screening, four high-performing antibody sequences were ultimately selected. By comparing various scoring metrics (such as pTM, ipTM, wpTM, and All pLDDT), the antibody sequence Antibody_82 was identified with the following scores: identifier score 0.644, pTM 0.462, ipTM 0.498, and wpTM as high as 83.094. In terms of plDDT, the All plDDT was 88.555, Binder plDDT was 80.558, and Target plDDT was 4.264. For the pAE metrics, Binder pAE was 11.062, Target pAE was 19.46, and Interaction pAE was 0.88. Furthermore, the Binder aligned lRMSD was 76.936. These data indicate that the antibody sequence shows excellent performance in terms of affinity and specificity.The amino acid sequence of Antibody_82 is as follows:

>Antibody_82_H EVQLVESGGGLVQPGGSLRLSCAASGFNIKDTYIHWVRQAPGKGLEWVARIY PTNGYTRYADSVKGRFTISADTSKNTAYLQMNSLRAEDTAVYYCSRYGYGG DYLFDYWGQGTLVTVSS >Antibody_82_L DIQMTQSPSSLSASVGDRVTITCRASQDVNTAVAWYQQQPGKAPKKLLIYSASFLYSGVPSRFSGSRSGTDFTLTISSLQPEDFATYYCQQHYTTPPTFGQGTKVEI K

Subsequently, we used the Humanness Prediction module to evaluate the humanization of Antibody_82. The prediction results showed that the humanization of Antibody_82 was 57.9% in Table 4, which is considered moderate, while its germline content was high, reaching 85.3%. Next, we performed druggability predictions using the Developability Prediction module. Antibody_82 showed a high purity of 95.887% in SEC-HPLC analysis, with a yield of 477.146 in Table 6, placing it at the high end of all tested samples. These data suggest that this antibody has a good prospect in drug development.

**Table 6.**
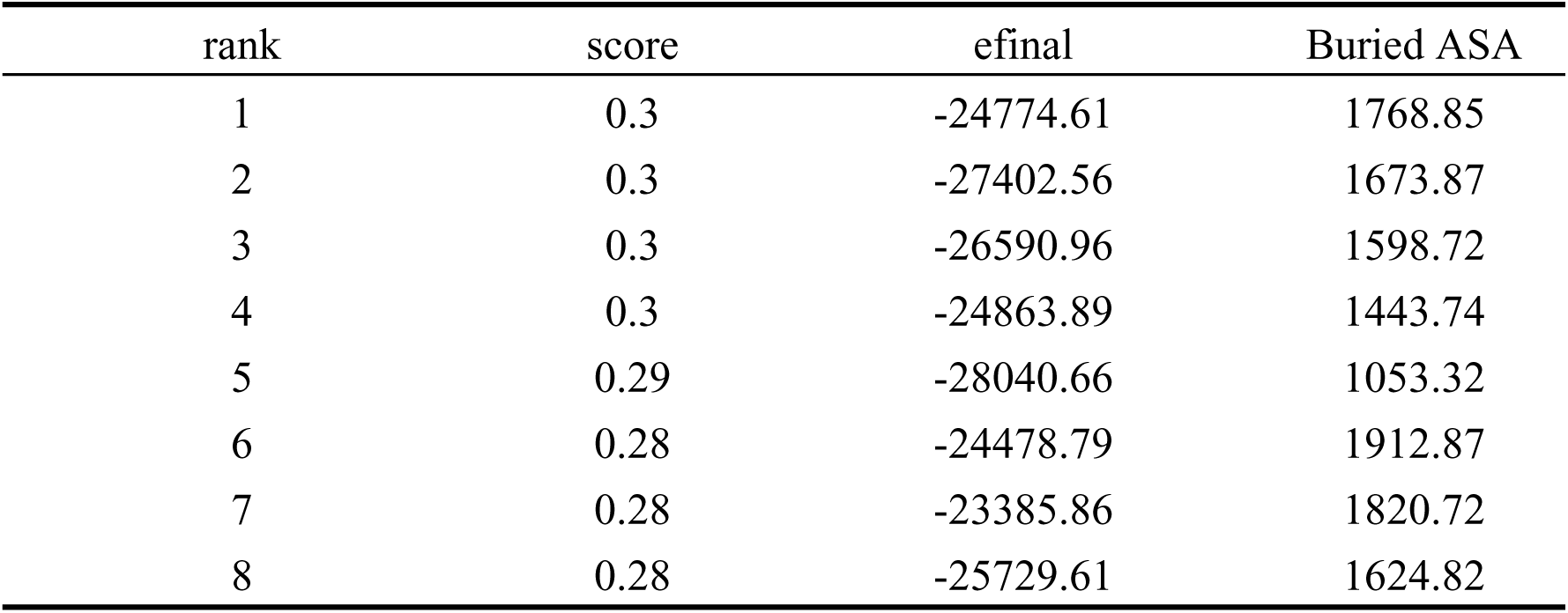
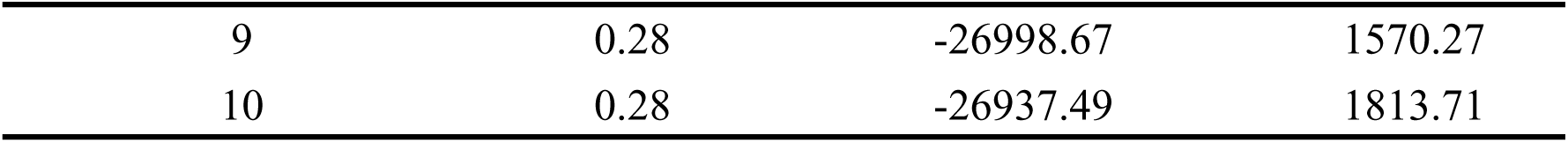
Ranking of Docking Poses Based on Scoring and Energetics.

### 3.10. Stability Analysis of Complex

To validate the binding effect of the LONP1 antigen with Antibody_82, we conducted an antigen-antibody docking experiment using the Antibody-Antigen Docking module. In the docking evaluation, we used the DockQ scoring system to assess the docking quality between the LONP1 antigen and the antibody. The DockQ scoring system is based on the similarity between the predicted and real structures and is widely used to evaluate the accuracy of protein-protein docking predictions.

Figure 16 shows the clustering heatmap of the predicted structures, which demonstrates a high degree of similarity between the predicted structures, and they closely resemble the real structure during the docking process.

**Figure 16.**
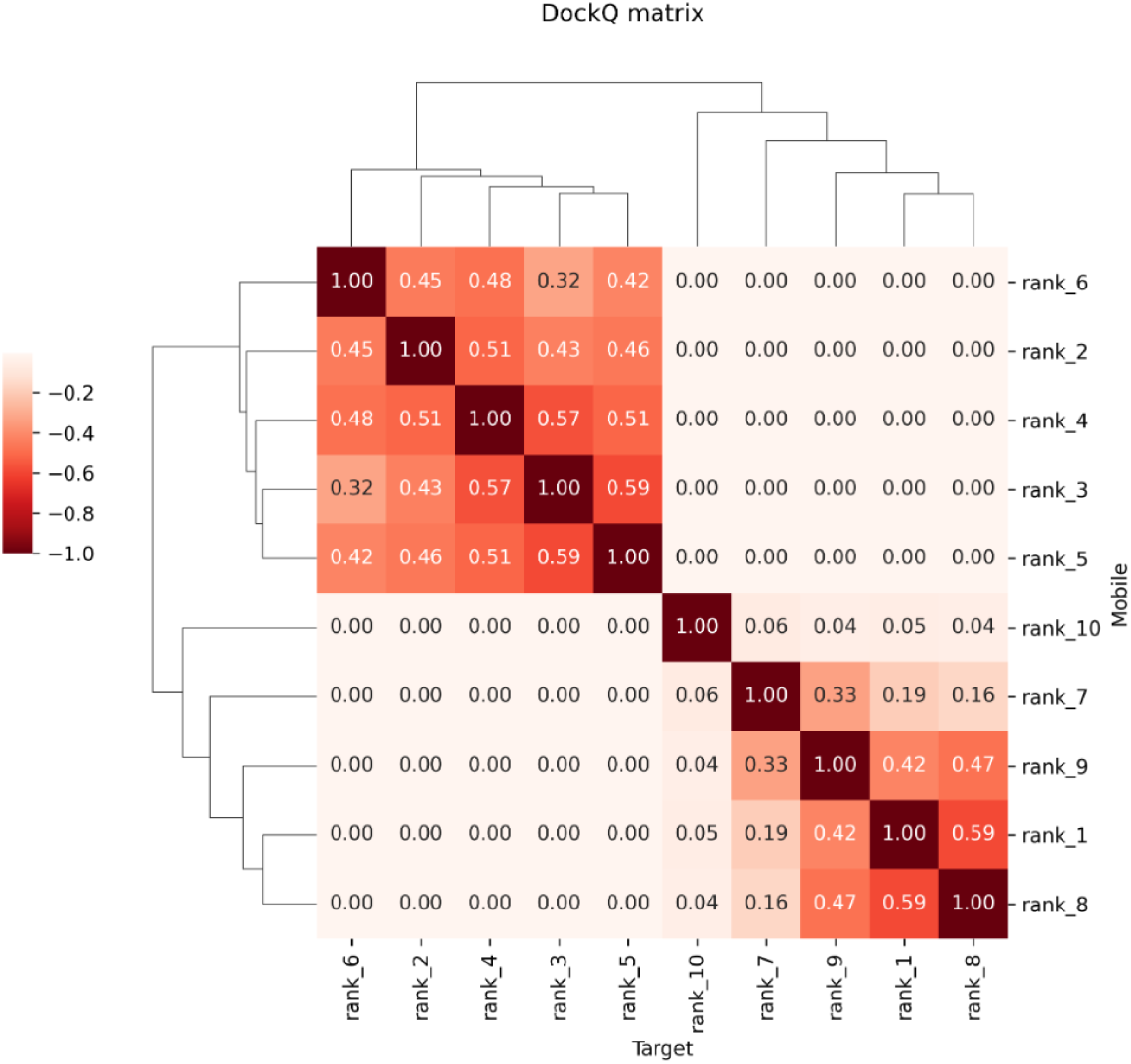
Clustermap of DockQ scores for predicted structures.

Table 6 provides the DockQ scores, final energy (efinal), and buried surface area (Buried ASA) for each docking structure. The DockQ score is an important indicator of docking quality; a higher score indicates that the predicted structure is more similar to the real structure. Final energy reflects the system’s energy during the docking process, with lower energy typically indicating a more stable docking structure. From the data in the table, the average DockQ score of the predicted structures was 0.29, and the average final energy was −25801.87, suggesting that the predicted structures are stable during docking.

To further validate the accuracy, reliability, and scientific soundness of the antigen-antibody complex, we conducted an antigen-antibody docking experiment using the GRAMM online docking platform and performed subsequent data analysis using PDBePISA.

Table 7 presents detailed information about the docking interface. ΔG (change in free energy, in kcal/mol) is a key thermodynamic parameter used to evaluate the stability of protein-protein docking. A lower ΔG value indicates a more stable docking structure. The analysis results show that the average ΔG value of the top five docking structures is −14.0 kcal/mol, suggesting that these docking structures are relatively stable.

**Table 7.**
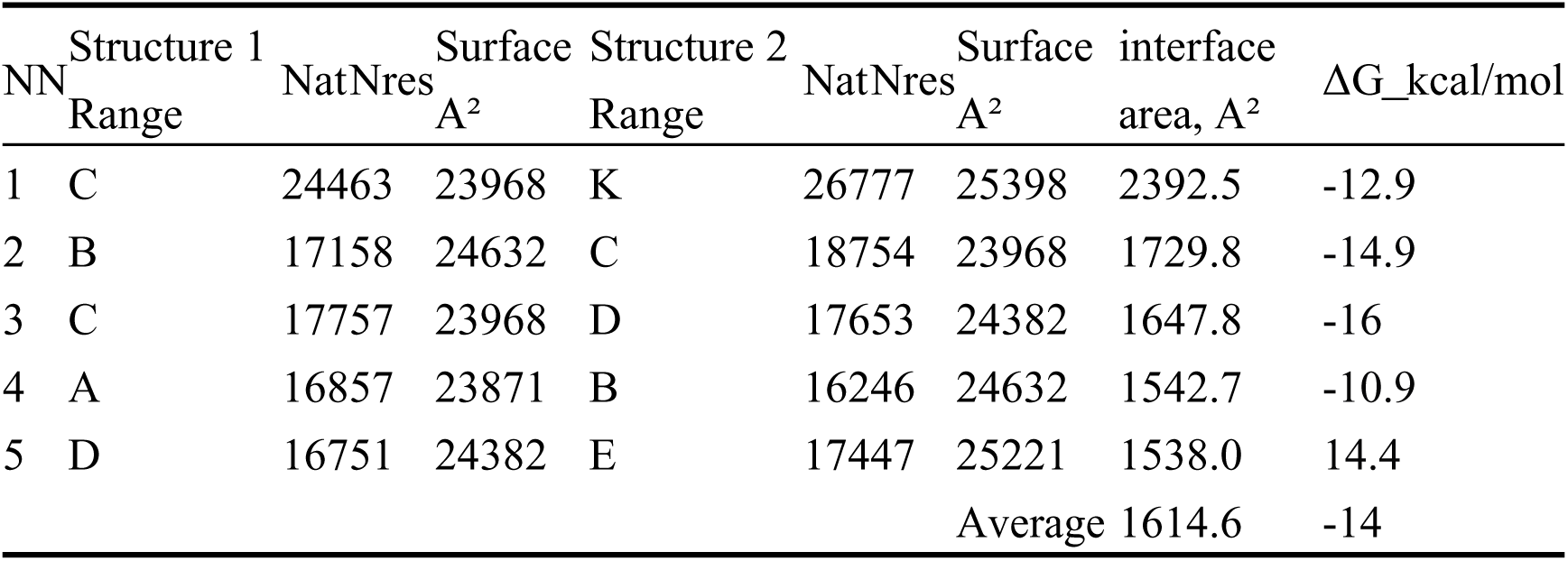
Summary of Docking Results for Complex Formation.

Further analysis revealed that the complementarity-determining regions (CDRs) of the LONP1 antigen and the Antibody_82 antibody formed multiple hydrogen bonds (Table 8) and salt bridge interactions(Table 9). The LONP1 antigen forms hydrogen bonds with several sites on the CDR of the Antibody_82 antibody, including interactions with amino acid residues such as GLU, SER, THR, TYR, VAL, ASN, ILE, LYS, and HIS. Additionally, the formation of salt bridges, particularly between amino acid residues like ARG and ASP, is crucial for stabilizing the complex structure. The average length of the hydrogen bonds is approximately 3.0 Å to 3.9 Å, indicating that these hydrogen bonds play an important role in the formation of the complex. The presence of salt bridges further enhances the stability of the binding, especially between amino acid residues with opposite charges. The interactions between these hydrogen bonds and salt bridges work together to promote the tight binding between the LONP1 antigen and the Antibody_82 antibody.

**Table 8.** The hydrogen bonds formed in the docking complex.

| ##Structure 1 | Dist. [A] | Structure 2 |
| --- | --- | --- |
| 1 C:TYR 492[OH] | 3.08 | K:GLU 388[OE2] |
| 2 C:GLN 677[N] | 2.47 | K:GLU 159[OE2] |
| 3 C:GLU 686[N] | 3.40 | K:THR 110[OG1] |
| 4 C:SER 687[N] | 3.35 | K:THR 110[OG1] |
| 5 C:SER 687[OG] | 3.15 | K:THR 110[OG1] |
| 6 C:SER 692[OG] | 3.40 | K:TYR 260[OH] |
| 7 C:VAL 737[N] | 3.81 | K:GLU 274[OE2] |
| 8 C:THR 738[OG1] | 3.10 | K:GLN 272[OE1] |
| 9 C:GLU 489[OE2] | 3.18 | K:ILE 206[N] |
| 10C:GLU 671[OE2] | 3.51 | K:ASN 268[ND2] |
| 11C:ARG 672[O] | 3.68 | K:ASN 201[ND2] |
| 12C:PRO 676[O] | 3.36 | K:SER 128[OG] |
| 13C:ALA 680[O] | 3.29 | K:SER 128[N] |
| 14C:ASP 685[OD2] | 2.91 | K:ARG 114[NH2] |
| 15C:ASP 685[OD2] | 3.40 | K:ARG 126[NH1] |
| 16C:GLU 686[OE2] | 2.71 | K:LYS 109[N] |
| 17C:GLU 686[OE2] | 3.27 | K:THR 110[N] |
| 18C:SER 687[OG] | 3.60 | K:ARG 126[NH1] |
| 19C:SER 692[O] | 3.83 | K:LYS 271[NZ] |
| 20C:SER 693[OG] | 2.35 | K:ASN 268[ND2] |
| 21C:GLU 736[O] | 2.52 | K:ARG 278[NH1] |
| 22C:GLU 736[OE2] | 3.09 | K:GLU 274[N] |
| 23C:GLU 736[OE2] | 3.76 | K:LYS 271[N] |
| 24C:TYR 921[O] | 3.18 | K:HIS 383[NE2] |

**Table 9.**
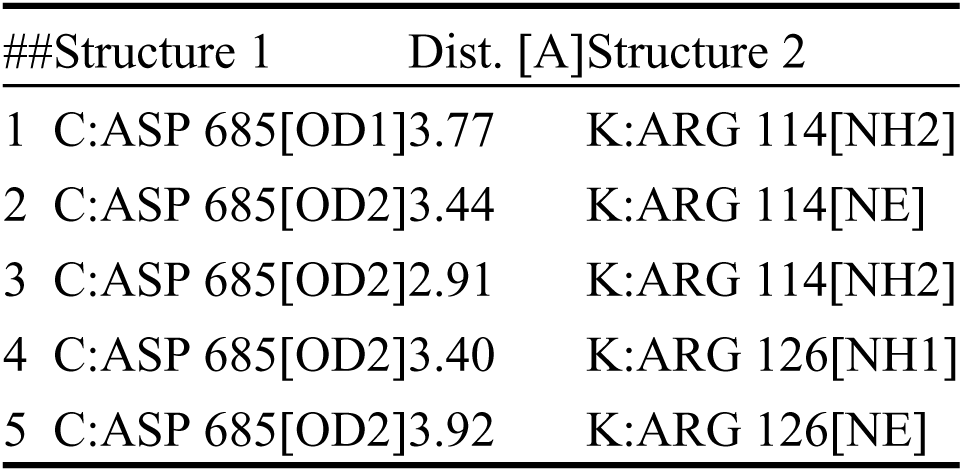
The salt bridges formed in the docking complex.

To further determine the stability of the complex, we conducted the following analyses using Gromacs:

Gibbs Free Energy Landscape (Figure 17): The chart displays the Gibbs free energy (ΔG) landscape from the antigen-antibody docking experiment, with energy values ranging from 2.5 kJ/mol to 15.0 kJ/mol, reflecting the stability of different conformations. Lower energy values indicate more stable docking conformations.

**Figure 17.**
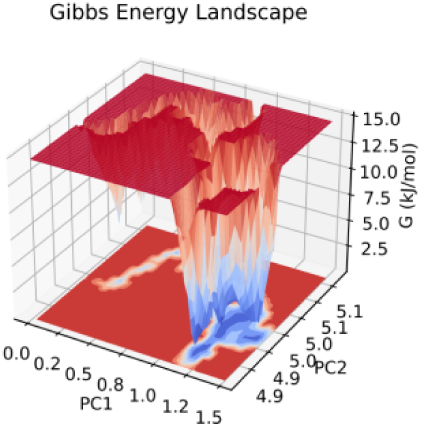
Gibbs Free Energy Landscape of Antigen-Antibody Docking Based on PCA.

RMSD (Root Mean Square Deviation) Chart (Figure 18): The chart shows the change in the RMSD of the protein backbone over time during the antigen-antibody docking experiment. Starting from zero, the RMSD value gradually increases and eventually fluctuates around 1.5 nm. This suggests that the protein structure underwent significant changes initially, then stabilized. The increase in RMSD may be related to conformational changes of the protein during the docking process, and the later stability indicates that the protein has found a stable docking conformation.

**Figure 18.**
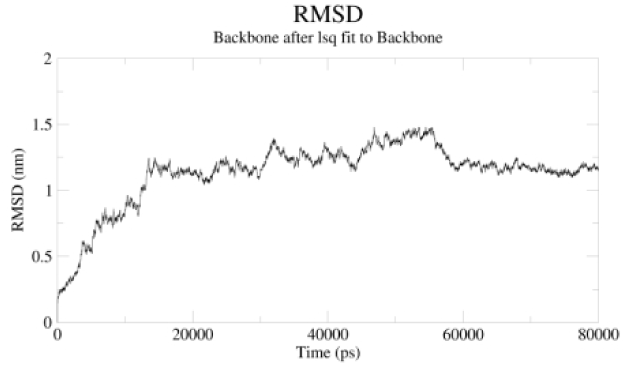
RMSD over time for protein backbone.

RMS Fluctuation Chart(Figure 19): This chart shows the RMS fluctuation of different residues of the protein during the docking process. Larger fluctuations were observed in regions with smaller residue numbers (likely the N-terminal), which are associated with flexibility or dynamic changes during the docking process. As the residue numbers increase, the fluctuations decrease, indicating that the C-terminal region is more stable.

**Figure 19.**
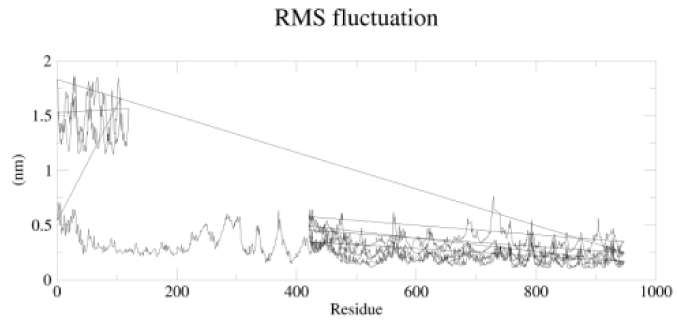
RMSD over time for protein backbone.

Radius of Gyration (Rg) Chart (Total and Axial Radii)(Figure 20): The chart shows that the total radius (Rg) of the protein molecule remained stable around 5 nm during the simulation, indicating the overall size stability of the protein molecule during the simulation.

**Figure 20.**
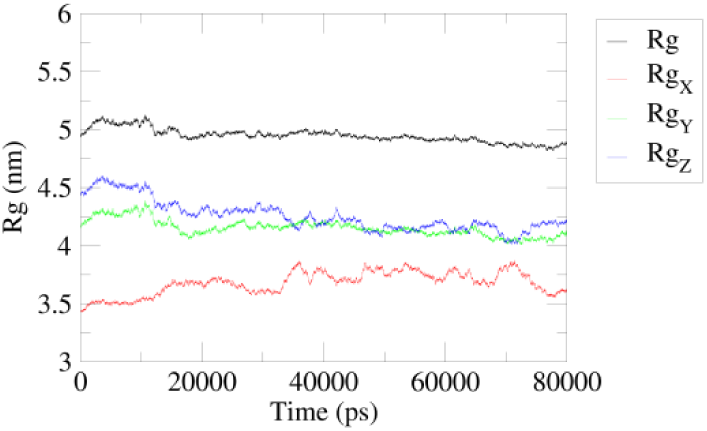
Radius of Gyration Over Time.

Additionally, we performed a visualization analysis using PyMOL. Figure 21 presents a visualization of Antibody_82, and Figure 22 shows the visualization of the interaction between the Lonp1 antigen and Antibody_82. The binding energy of the antigen-antibody complex is −12.9 kcal/mol(Table 10), further indicating the good structural stability of the complex.

**Figure 21.**
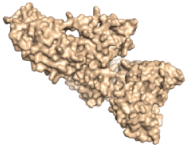
3D Molecular Structure Visualization of LONP1.

**Figure 22.**
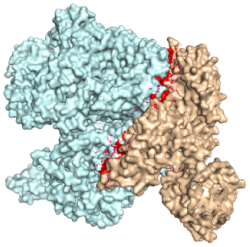
3D Structure Visualization of LONP1-Antibody82 Complex.

**Table 10.** Binding energy between LONP1 and Antibody_82.

| LONP1 | Antibody_82 | Binding Energy (kcal/mol) |
| --- | --- | --- |
| Chain C | Chain H | -12.9 |
| Chain B | Chain H | -5.1 |

Video 1 demonstrates the stable binding structure of the LONP1 antigen in complex with the Antibody_82 antibody.

**Vedio 1.**
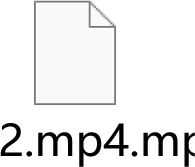
Dynamic Docking of LONP1-Antibody82 Complex.

The ATP-binding site of LONP1 is situated at positions 523-530 on the C-chain. Validation with PyMOL revealed that the binding region between the LONP1 antigen and Antibody_82 spans from positions 421 to 630 on the C-chain(Table 11), which includes the ATP-binding site. The findings demonstrate that Antibody_82 binds to LONP1 at a site that aligns with expectations, and the docking structure is stable. The antigen-antibody interaction is capable of disrupting the ATP-binding site of LONP1, potentially inhibiting its function. Overall, Antibody_82 exhibits good structural stability and presents favorable prospects for pharmaceutical development.

**Table 11.** Binding site between LONP1 and Antibody_82.

| LONP1-binding antibody | Binding site |
| --- | --- |
| Chain C | 421-630 |
| Chain B | 480-740 |
| Chain D | 720-726 |

## 4. Discussion

The trend analysis indicates that the number of publications in the fields of prostate cancer and mitochondria has significantly increased in recent years, particularly between 2015 and 2017. This growth trend reflects the rapid development of these fields and the increasing research interest they are attracting. The underlying reasons for this rise may include the following factors: First, the aging population has led to an increase in the incidence of prostate cancer, which has subsequently generated more research attention. Second, advances in biomolecular research technologies have enabled researchers to investigate more deeply into the role of mitochondria in prostate cancer. Additionally, the growing public awareness of prostate cancer has increased the demand and attention for research in this area[22]. The United States is a leader in this field, likely due to ample research funding, advanced research infrastructure, and a concentration of scientific talent. Since 2020, China has seen a rise in annual publications, reaching the top spot globally, which is closely related to the country’s increasingly aging population and the promotion of cancer early screening.

Additionally, the articles “Guidelines for the use and interpretation of assays for monitoring autophagy (3rd edition)” published by Daniel J. Klionsky in 2016 and “Guidelines for the use and interpretation of assays for monitoring autophagy (4th edition)” published in 2020 have become the most cited works in this field. These two papers delve deeply into the process of autophagy, with particular focus on its mechanisms in mitochondria and prostate cancer.

Autophagy is an intracellular process that involves the formation of a double-membrane structure called the autophagosome, which encloses cellular components that need to be degraded, including damaged proteins and organelles such as mitochondria. This process is essential for maintaining cellular health and plays a crucial role in responding to various disease states.

Mitochondrial autophagy, the process of degrading damaged or unnecessary mitochondria, plays a critical role in the development and therapeutic response of prostate cancer. By eliminating damaged mitochondria, mitochondrial autophagy reduces oxidative stress and DNA damage, which helps inhibit tumor progression. This process is also closely linked to cell survival and death. In some cases, it promotes apoptosis of tumor cells, while in others, it can suppress cell death, thereby influencing the survival of tumor cells.

Furthermore, mitochondrial autophagy alters the metabolic and signaling molecules within the tumor microenvironment, affecting tumor proliferation, migration, and metastasis. It also plays a potential role in immune evasion and immune surveillance by modulating immune cells. Additionally, mitochondrial autophagy influences angiogenesis and the function of tumor stromal cells, which contribute to tumor growth and metastasis[23–25].Mitochondrial autophagy can also regulate drug sensitivity. Certain anticancer drugs enhance their antitumor effects by inducing mitochondrial autophagy, helping to overcome drug resistance[26].

Mitochondria have become a focal point in prostate cancer research, with recent studies highlighting issues such as drug resistance and radioresistance. For patients with advanced prostate cancer, androgen deprivation therapy (ADT) is the primary treatment. However, nearly all patients undergoing ADT eventually develop drug resistance within 18 to 36 months, leading to the progression to CRPC(castration-resistant prostate cancer)[27].CRPC is an advanced form of prostate cancer characterized by the ability of cancer cells to continue growing and spreading despite androgen deprivation. The prognosis for CRPC patients is poor, with an average survival time of only 9 to 13 months[28]. Recently, understanding the mechanisms behind CRPC development and exploring therapeutic strategies has become a major research focus.Treatment options for CRPC primarily include:New androgen receptor pathway inhibitors such as enzalutamide and abiraterone, which slow down cancer cell growth by inhibiting androgen signaling[29]. Chemotherapy drugs like docetaxel and cabazitaxel, which are used to suppress cancer cell division. Immunotherapy, including treatments like Sipuleucel-T and immune checkpoint inhibitors (e.g., PD-1/PD-L1 inhibitors). Radiation therapy, which can help alleviate pain caused by bone metastasis. Targeted therapies such as PARP inhibitors (e.g., olaparib) that target cancer cells with DNA repair deficiencies. Bone metastasis treatments, including castration therapy and bone-modifying agents like bisphosphonates and denosumab. Radionuclide therapy, such as Radium-223, which offers treatment options for patients with bone metastases. The search for new therapies continues, as drug resistance and the complex mechanisms of CRPC present significant challenges in the fight against prostate cancer[30,31].

Despite achieving some clinical success, the treatment of CRPC still faces many challenges, with resistance being one of the biggest issues. Even when patients initially respond to treatments such as androgen receptor inhibitors, tumors often develop resistance over time through new escape mechanisms. Additionally, many treatments, especially chemotherapy and immunotherapy, can cause significant side effects that severely impact patients’ quality of life. The variability in how different patients respond to the same treatment makes predicting treatment outcomes even more complex. For patients with advanced CRPC, particularly those who have undergone multiple lines of therapy with continued disease progression, treatment options are extremely limited[32].

The study of resistance mechanisms remains a critical direction in the current field of prostate cancer research. Researchers are dedicated to developing new treatment strategies, and targeted intervention on mitochondria is a promising approach, aiming to enhance the effects of existing therapeutic drugs by modulating mitochondrial function. Although LONP1 is a potential therapeutic target, there has been limited research and development on inhibitory antibodies against it, with no therapeutic antibodies targeting LONP1 available on the market. Therefore, through the GeoBiologics platform, we virtually designed and screened specific antibodies against LONP1 to hinder its function, hoping to treat CRPC by targeting and inhibiting mitochondrial function.

However, delivering targeted antibodies into cells remains a challenge in the field of biomedical research. Antibodies typically cannot directly penetrate the cell membrane, so various strategies and techniques are required to facilitate their entry into cells. Common methods include:Cell-penetrating peptides (CPPs), such as TAT peptides, which are commonly used to enhance the internalization of antibodies or other biomolecules by binding to them.Nanocarrier systems, which use nanoparticles (such as lipid nanoparticles or polymer nanoparticles) to encapsulate the antibody and facilitate its entry into cells through endocytosis.Viral vectors, such as adenoviruses or lentiviruses, can be engineered to deliver antibody genes or antibody proteins. These vectors enable the transfer of antibody genes or proteins into cells for expression or action.Ultrasound-mediated delivery, which utilizes the physical effects of ultrasound by applying rapid pressure fluctuations to the cell membrane, creating temporary reversible holes, thus allowing antibodies to enter the cells[33–36].These methods help deliver targeted antibodies into cells through various mechanisms, thereby enhancing treatment efficacy.

Targeted antibody combination therapy has shown significant potential in the treatment of prostate cancer, especially in advanced stages such as metastatic castration-resistant prostate cancer (CRPC). Targeted antibody combination therapy can enhance treatment efficacy from multiple directions, overcome resistance, and reduce side effects, ultimately improving patients’ survival and quality of life[37].

In our bibliometric analysis,, we identified 152 studies on mitochondrial-targeted therapy for prostate cancer, with 29 studies focusing on targeted and combination therapies. The goal of these therapies is to enhance efficacy through multiple mechanisms and overcome the limitations of single treatments. Common targeted antibody combination therapies include combinations with hormonal therapies (Abiraterone or Enzalutamide), chemotherapy (Docetaxel), radiation therapy (EBRT), anti-angiogenesis therapies (Olaparib, Anlotinib), bone metastasis treatment (Denosumab), and PARP inhibitors (Olaparib)[38–41]. In clinical practice, the appropriate combination therapy strategy is chosen based on the patient’s specific condition and the molecular characteristics of the cancer.

## 5. Conclusion

The research mentioned above provides new perspectives for the treatment of CRPC. Although mitochondrial-targeted therapy shows promising potential in CRPC treatment, fully realizing its potential requires overcoming existing challenges through rigorous scientific research and clinical trials. With the continuous advancement of science and technology, there may be more groundbreaking discoveries in the future, offering more effective and safer treatment options for CRPC patients.

## Supporting information

vedio

## 6. Acknowledge

This work was supported by awards from the Natural Science Foundation of Jiangsu Province (BK20231189), and PanFeng Innovative Team Project of the The Third Affiliated Hospital of Soochow University.

## Notes

### Competing Interest Statement

The authors have declared no competing interest.

